# Larval brooding damselfishes and shifting body proportions: does pelagic larval swimming constrain reef fish morphology?

**DOI:** 10.1101/2023.09.25.559426

**Authors:** J.P. Lyons, K.D. Kavanagh

## Abstract

The vast majority of reef fishes, including damselfishes, have a pelagic larval stage that ends when any surviving larvae swim to a reef to settle. An extremely rare alternative lifestyle is ‘larval brooding’, where both parents protect larvae on the reef for months until they disperse nearby. The monophyletic clade of larval brooders includes two genera *Acanthochromis* and *Altrichthys*. In examination of the skeletons of these damselfishes, we found that all species of damselfish that brood larvae on the reef have a uniquely increased precaudal vertebral count, while all typical damselfishes have an invariable vertebral count with a greater proportion of caudal vertebrae. To explore the significance of the vertebral differences, we measured body proportions of larval brooders vs relatives with the typical pelagic larval stage. We found increased body cavity area and reduced muscle area in the larval brooder *Acanthochromis polyacanthus*. Furthermore, populations of *Acanthochromis* throughout its range have evolved significantly different proportions. In a comparison of known larval swimming ability among reef fishes, the larval brooders performed most poorly. We propose that when larval brooding evolved, relaxed selection on larval swimming performance allowed a shift in body proportions to favor a larger body cavity and altered axial patterning. Enlarged body cavity gives a fitness advantage as females could hold more of the large eggs and increase clutch size.

## Introduction

Marine fishes typically invest very little in individual offspring, spawning 100,000s or millions of fertilized eggs over a year with no parental care or only brief demersal egg care (Leis and McCormick, 2002; Thresher, 1984). The adaptive significance of this “r-selected” (Macarthur and Wilson, 1967) life history as a strategy to avoid heavy predation on the reefs is well supported by empirical and theoretical studies (Mihalitsis et al. 2022; Dytham and Simpson 2007; Johannes, 1978).

Pelagic eggs and larvae are initially fully planktonic, dispersing with the currents, but as they grow over weeks, the (very few) surviving larvae gain in size, sensory competence, and swimming ability (Leis and McCormick, 2002). It is estimated that less than 0.001% of pelagic larvae survive to settlement (∼20% loss per day; Houde and Zastrow, 1993), but even that small number is greater than survival on the reef as a larva, which is essentially zero without biparental care (Nakazono, 1993). Such intense predation pressure may have driven the evolution of early life stages moving to the open ocean until they reach older juvenile stages (Dytham and Simpson, 2007).

Only a very few out of ∼20,000 marine fishes lack the pelagic larval stage (Leis, 1991). An extremely rare life history is ‘larval brooding’, where both parents protect larvae on the reef for months until they disperse nearby. Larval brooding evolved in one small clade of the damselfishes (Pomacentridae), that includes two genera *Acanthochromis* and *Altrichthys*. This life history has eliminated the pelagic larval stage. These four related damselfish species are known to brood their young on the reef after hatching, somehow overcoming the “ecological constraint” of heavy larval predation on the reef. Especially well known is the most widespread of these species *Acanthochromis polyacanthus* (a monotypic genus), which has prolonged bi-parental care of the brood, staying together as a family unit for 2-3 months (Kavanagh, 2000; Robertson 1973). This high level of parental investment is countered by an annual fecundity for *Acanthochromis* several orders of magnitude lower than other reef fishes (Fig. 1), indicating a major evolutionary shift toward “K-strategy”. This shift is correlated with many unique morphological, behavioral, and life history traits (Table 1) and is extremely rare in other fish taxa (Allen 1999). The reproductive biology and early life history of *Altrichthys* spp. are not known.

**Table 1.**
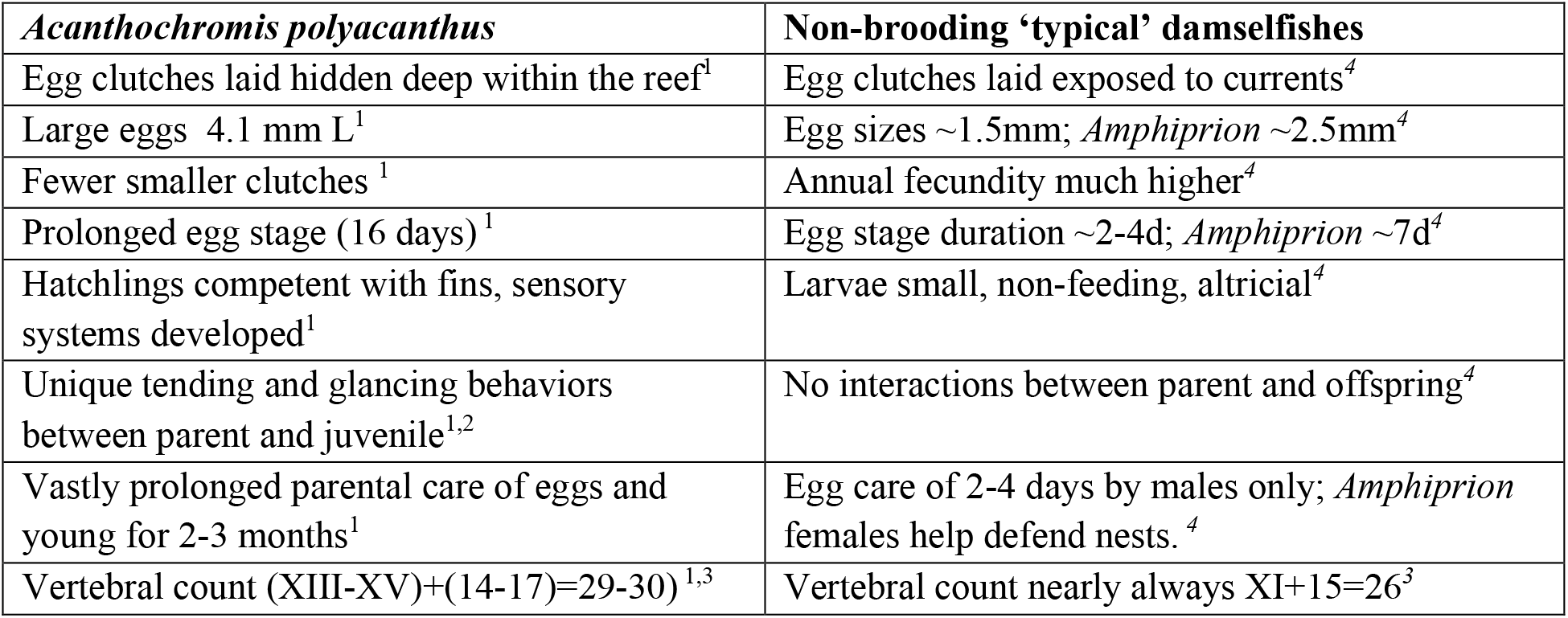
Major phenotypic differences shown by Acanthochromis polyacanthus compared with the other non-brooding damselfishes. (^1^Kavanagh, 2000; ^2^Kavanagh, 1998; ^3^Kavanagh and Leis, 2003; ^4^Frédérich, B., & Parmentier, E. 2016).

| *Acanthochromis polyacanthus* | Non-brooding ‘typical’ damselfishes |
| --- | --- |
| Egg clutches laid hidden deep within the reef<sup>1</sup> | Egg clutches laid exposed to currents<sup>4</sup> |
| Large eggs 4.1 mm L<sup>1</sup> | Egg sizes ~1.5mm; *Amphiprion* ~2.5mm<sup>4</sup> |
| Fewer smaller clutches<sup>1</sup> | Annual fecundity much higher<sup>4</sup> |
| Prolonged egg stage (16 days)<sup>1</sup> | Egg stage duration ~2-4d; *Amphiprion* ~7d<sup>4</sup> |
| Hatchlings competent with fins, sensory systems developed<sup>1</sup> | Larvae small, non-feeding, altricial<sup>4</sup> |
| Unique tending and glancing behaviors between parent and juvenile<sup>1,2</sup> | No interactions between parent and offspring<sup>4</sup> |
| Vastly prolonged parental care of eggs and young for 2-3 months<sup>1</sup> | Egg care of 2-4 days by males only; *Amphiprion* females help defend nests.<sup>4</sup> |
| Vertebral count (XIII-XV)+(14-17)=29-30)<sup>1,3</sup> | Vertebral count nearly always XI+15=26<sup>3</sup> |

**Figure 1.**
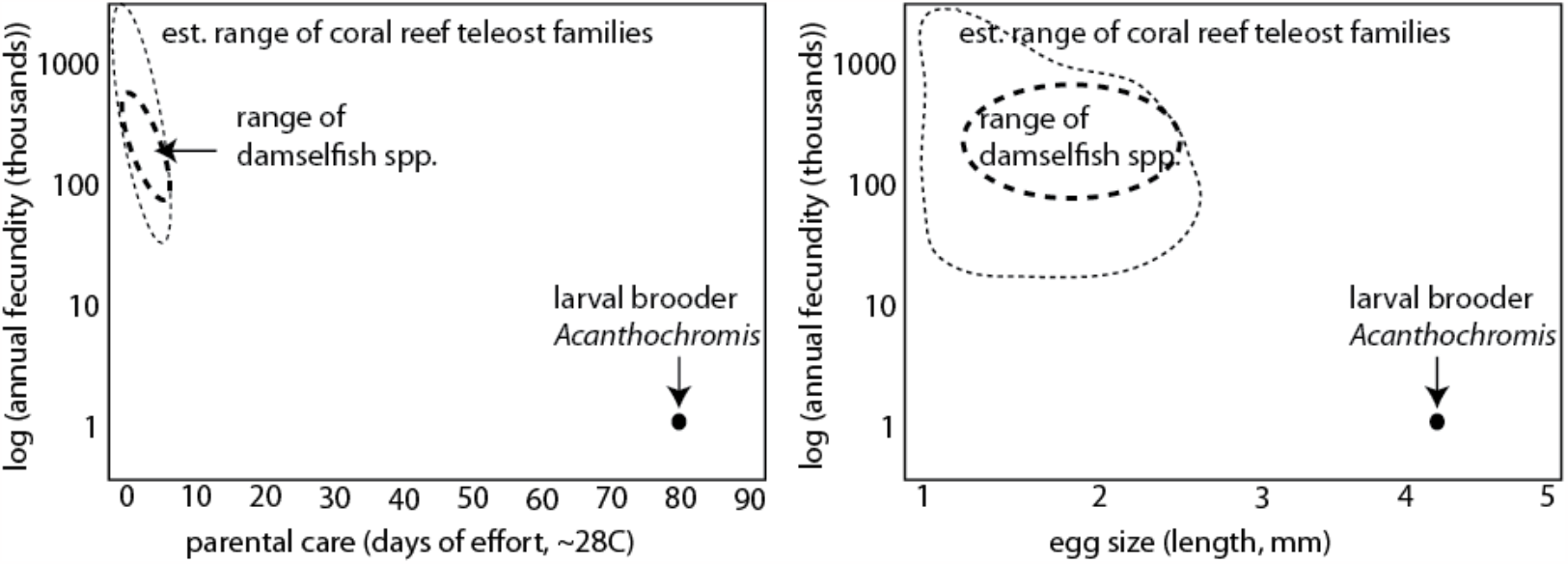
Reproductive investment per offspring is dramatically higher in <u>Acanthochromis</u>, and its estimated annual fecundity per female is orders of magnitude lower, than other damselfishes (data from Thresher 1984, Kavanagh 2000, and Kavanagh and Alford, 2003).

The pelagic stage ends when the older larvae orient using multiple sensory cues (Lecchini et al., 2005; Leis and McCormick, 2002) and swim to a reef to settle. In recent years, the capability of pelagic fish larvae to orient and swim for long sustained periods (Fisher and Leis 2010; Stobutzki and Bellwood 1997) has become clear.

*Acanthochromis* lacks the pelagic stage entirely, and the species shows many derived reproductive, morphological, behavioral, and life history correlates with larval brooding on coral reefs (Table 1; Figure 2; (Kavanagh, 2000; Kavanagh and Alford, 2003). One particularly striking phenotypic divergence is the vertebral count – *Acanthochromis* has typically 3-4 more precaudal vertebrae than confamilial species.

**Figure 2.**
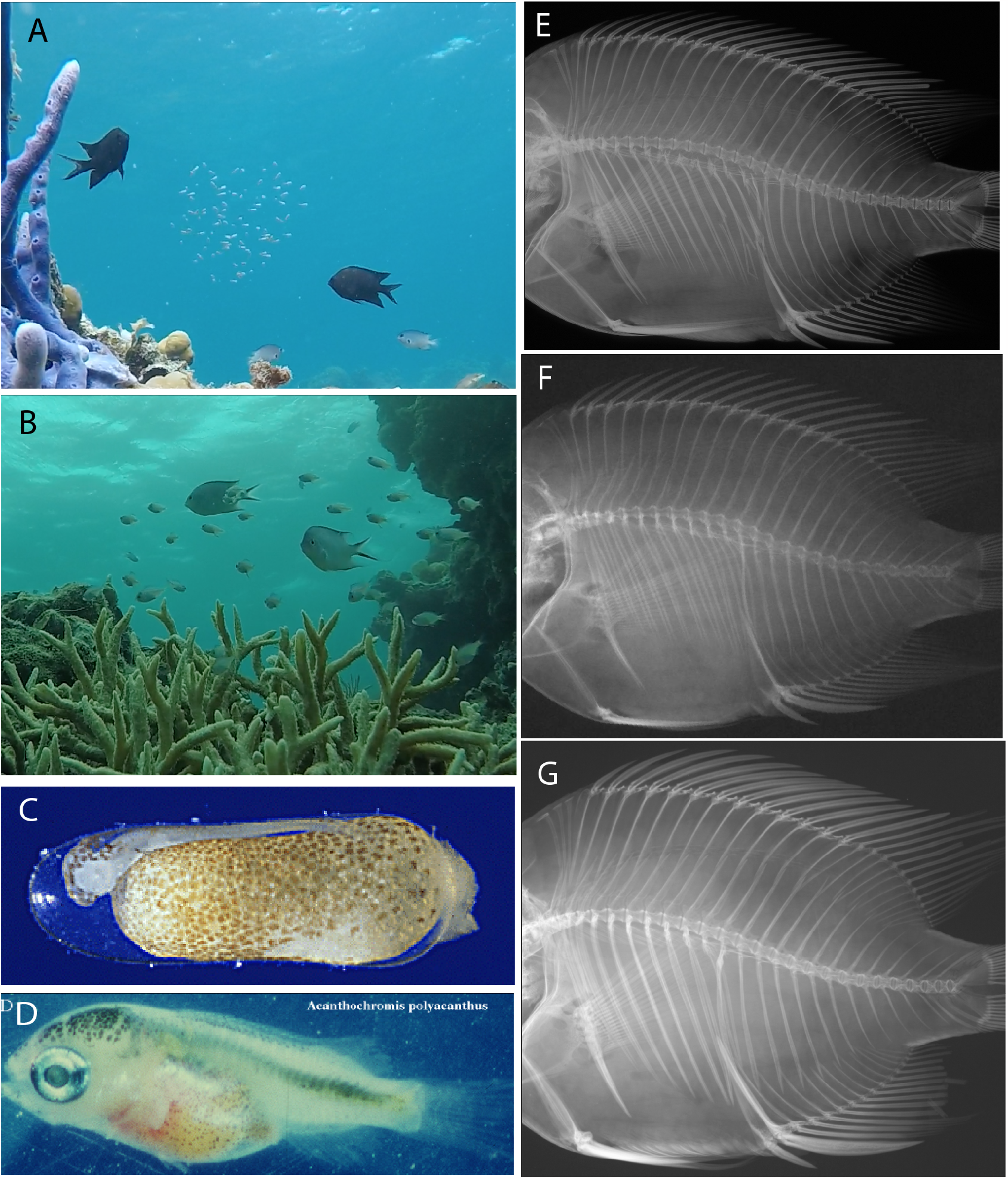
Ontogeny, morphology and parental care behavior of brooding damselfishes. A. Extended biparental care: Acanthochromis polyacanthus family with a young brood and B. Altrichthys azurelineatus family with an older brood (both from Busuanga, Philippines). C. Acanthochromis embryo at 2 days after fertilization (4mm); D. Acanthochromis hatchling (16 days after fertilization; 5.5mm); E.-G. Radiographs of Acanthochromis (XV+14=29) (E), Altrichthys curatus (XII+15=27) (F), and Amblyglyphididon curacao (XI+15=26) (G) showing expanded precaudal vertebral counts in larval brooders.

We suspected that the difference in meristic counts may be related to the life history shift to larval brooding. All ∼350 species of damselfishes are reported to have an invariable vertebral count of XI+15=26 (11 precaudal, 15 caudal, 26 total vertebrae, with rare variants), with the sole exception of *Acanthochromis*, which has 3-4 additional vertebrae concentrated in the precaudal region (XIII-XV+14-17=29-30) and greater variability among individuals (Kavanagh et al., 2000; Kavanagh and Leis, 2003; this study). We wondered whether this rare precaudal vertebral increase facilitated the evolution of brooding by allowing the body cavity to hold more of the relatively huge eggs, increasing reproductive output in the risky reef environment (Fig 1, Table 1). We tested this idea by comparing regional body proportions and vertebral counts in larval brooders and non-brooders, including the first meristics and morphometric analysis of *Altrichthys* spp., a little-known larval brooding genus which is the sister clade of *Acanthochromis* (Bernardi, 2011) and is endemic to northern Palawan and the Calamanian Islands.

## Methods

### Specimens

Lateral radiographs of 620 subadult or adult damselfish of 10 genera and 51 species were obtained from the American Museum of Natural History, the Australian Museum, the Western Australia Museum, the Bernice P. Bishop Museum, and the Museum of Comparative Zoology. (Suppl. Info. 1). The two larval brooding genera *Acanthochromis* and *Altrichthys w*ere compared with the phylogenetically closest genera that had the typical pelagic larval stage *Hemiglyphidodon, Amblyglyphidodon*, and *Neoglyphidodon*, as well as selected damselfish genera that were more phylogenetically distant and with morphologically diverse body shapes (*Amphiprion, Dascyllus, Chromis, Lepidozygous*, and *Pomacentru*s). *Acanthochromis* specimens originated from several locations, so we conducted further examination of morphometric divergence by populations, including Philippines, Papua New Guinea, Solomons Islands, Vanuatu, and southern Great Barrier Reef (Suppl. Info. 1).

### Vertebral counts

Precaudal vertebrae are determined by presence of neural spines and paired ribs. Caudal vertebrae are identified by having a single hemal arch and neural spine. The insertion of the anal spine marks the transition between precaudal and caudal region. Ranges and frequencies of precaudal and caudal vertebral counts for each species were recorded.

### Precaudal vertebral expansion

To test how where the vertebral column changes as body cavity expands, we counted the number of vertebrae in anterior, middle, and posterior regions of the precaudal straight-line A-P distance. We counted the precaudal vertebral regions for five specimens from each of the larval brooding genera *Acanthochromis* and *Altrichthys*, as well as a close non-brooding relative *Amblyglyphidodon* and two non-brooding relatives with unusual body shapes, the rounded *Dascyllus* and the elongate *Lepidozygous*. Vertebrae in these regions were counted, and vertebrae spanning two regions were visually estimated in tenths of a vertebra. Mean numbers of anterior, medial, and posterior precaudal vertebrae for each genus were tested, using a Welch One Way ANOVA with post hoc Games-Howell multiple comparisons for differences between individual genera.

### Morphometrics

Body proportions and lengths were measured on each specimen using ImageJ tools (https://imagej.nih.gov). Standard Length (SL) was measured from a line was drawn from the anterior tip of each fish’s jaw to the posterior end of the caudal peduncle. Two dimensional area from the lateral view was divided into head, body cavity, and total body area, as defined below. Muscle area was calculated as total body area -(head area and body cavity area). The head region of damselfish in this study was defined as a polygon defined by enclosing the skull and areas anterior to the first neural spine and insertion of the pelvic fins. The body cavity region is defined as the space within the polygon drawn by placing a point at the anterior end of each precaudal vertebra, the insertion of the pelvic fin, and the insertion of the anal fin. Total body area of each fish in 2D is defined as all fish tissue that is not a fin ray. A polygon was drawn along the periphery of the entire fish body including the caudal peduncle but excluding the fins. By subtracting the area of the Head and Body Cavity regions from Total Body Area, the Muscle Area was computed.

### Statistical comparisons of body regions

For comparisons, we first grouped individuals in our sample by genus, creating 10 groups with two categories: two larval brooding genera and eight non-brooding genera. *Acanthochromis* measurements, which were particularly variable, were later divided into populations by geographic location: Southern Great Barrier Reef, Papua New Guinea, Solomon Islands, Vanuatu, and Philippines for further analysis. Means for all measured anatomical regions were tested by Welch’s One Way ANOVA with post hoc Games-Howell multiple comparisons. Raw area measurements from ImageJ were converted into proportions. Because relative head areas were found to differ among species, we subtracted head area from total body area, thus postcranial body area (Total Body Area – Head Area) was the unit for normalizing each individual’s measurements of body regions. The area measurements of interest were thus only two size-standardized and shape-independent postcranial proportions – body cavity area and muscle area. Type III ANOVAs (Welch test) with Games-Howell pairwise post hoc tests were performed using JASP (Version 0.10.2.0) to test differences in body cavity area and muscle area among taxon.

### Larval swimming published data

To compare larval swimming ability among species, we used data compiled in a recent review that included Ucrit swimming speed data from both young brooded larvae of *Acanthochromis* and late larval stages of 10 non-brooding damselfish genera (Fisher et al., 2022), which were approximately similar in age and/or size. Fish in the dataset were pooled to the genus level.

## Results

### Vertebral counts

Compared with all non-brooding (“typical”) damselfishes, we found that all the brooding damselfish species had a higher precaudal vertebral count. *Acanthochromis polyacanthus* had 3-4 more precaudal vertebrae (69% (87/126) had 14, 29% (37/126) had 15, and 1.6% (2/126) had 13), and a caudal vertebral count that was similar, but more variable, than non-brooders (95% (120/126) had 15; 5% had 14) (Suppl. Info. 2). *Altrichthys azurelineatus* and *Altrichthys curatus* both had one additional precaudal vertebra (12; 1/18 had 11) and, again, a similar number of caudal vertebrae as non-brooders (15). Consistent with published literature, all non-brooding damselfishes have an invariable vertebral count of XI+15=26. The only exception was *Hemiglyphidodon a*nd *Lepidozygous:* each had a single individual with 10 precaudal vertebrae instead of the typical 11. (Suppl. Info. 1)

### Precaudal expansion in Acanthochromis and other genera

To examine the pattern of precaudal vertebrae segmentation, we measured the relative size of precaudal vertebrae of *Acanthochromis*. The average size of precaudal vertebrae was smaller in *Acanthochromis* compared with other genera (Suppl. Info.) We also assessed what part of the precaudal region the additional vertebrae were found by counting anterior, middle, and posterior thirds of the precaudal vertebral column. The additional vertebrae of *Acanthochromis* are distributed across the precaudal region ‘thirds’ with approximately 1 more per section (Table 2). *Altrichthys* has one more precaudal vertebra, and the addition was distributed across the precaudal region, with about 1/3 of a vertebra more per section. In contrast, *Lepidozygous tapeinosoma* is a non-brooder with an unusually elongate body shape and had larger anterior and smaller posterior vertebrae (Table 2).

**Table 2.**
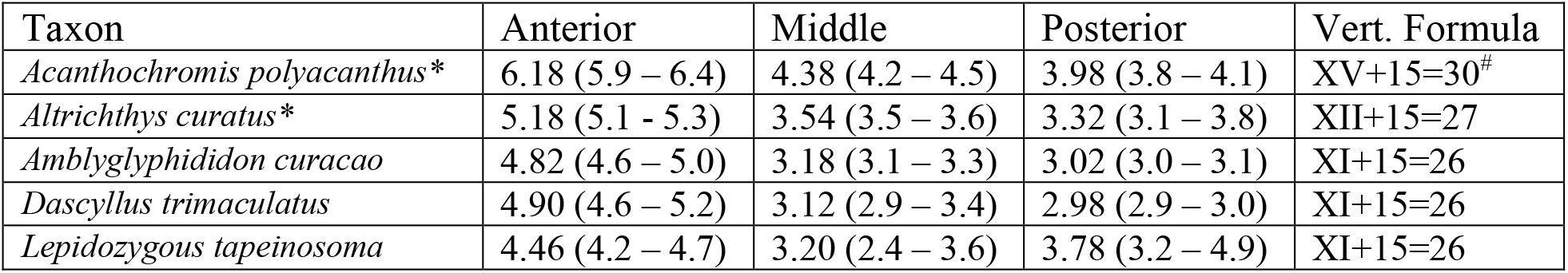
Number of vertebrae in each third of the precaudal vertebral section (mean and range in parentheses). Vertebral patterning differs among taxa. More vertebrae in a section means that vertebrae are smaller; fewer vertebrae/section mean larger vertebrae. N=5 for each taxon. ^*^larval brooder; ^#^most common phenotype. The additional vertebrae of Acanthochromis and Altrichthys are distributed across the precaudal region ‘thirds’. Lepidozygous tapeinosoma has an unusually elongate body shape and has larger anterior and smaller posterior vertebrae.

### Body proportions

To explore the consequence of the increased vertebral number in the brooding damselfishes, we measured body proportions of larval brooders vs non-brooding relatives. We hypothesized that additional precaudal vertebrae would lead to greater relative body cavity area, standardized to total postcranial 2D body area (from radiographs), in larval brooders. Because head area varied among species, head area was subtracted from total body area to compare more equivalently the postcranial proportions between body cavity and muscle area among taxa. Levene’s test for equality of variances found significant differences among genera, therefore we used Welch’s ANOVA to test for significant differences in body proportions among taxa.

We found that larval brooders had a significantly higher body cavity (=abdominal) area (Welch’s ANOVA: F=73; df=9; p>0.001; Fig. 3). Games-Howell post hoc tests show that *Acanthochromis* had significantly greater body cavity area than all other genera except *Pomacentrus*. Body cavity area of *Altrichthys* spp. (the other larval brooding genus with one additional precaudal vertebra) was significantly greater than 5/8 non-brooding species (Suppl Info. 2). Body cavity area was smallest in *Lepidozygous* and *Amphiprion (*Fig. 3).

**Figure 3.**
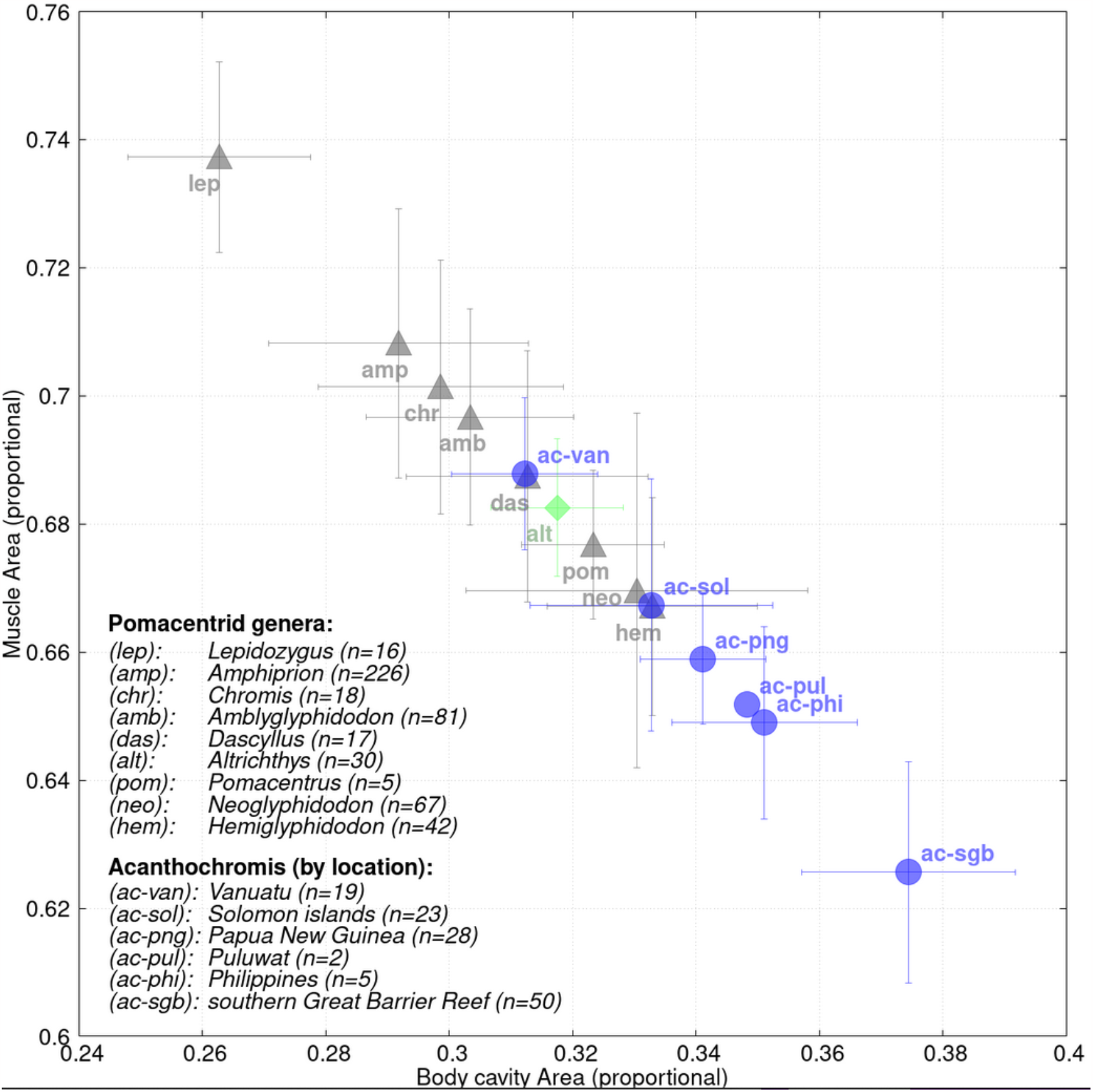
Body cavity and muscle area by genus. Symbols are means and lines are one standard deviation. Acanthochromis populations are divided by location. Acanthochromis and Altrichthys are larval brooders and lack the pelagic larval stage.

We further examined whether the expanded body cavity area in *Acanthochromis* was due to precaudal body length, body depth, or a combination. Precaudal length (i.e. body cavity A-P length) was significantly different among genera. Games-Howell post hoc comparisons by genus showed that precaudal length was greater in *Acanthochromis* than other genera except for *Pomacentrus* (Suppl. Info. 2). The larval brooders *Altrichthys* precaudal length was significantly greater than about half of the other genera. Precaudal length was smallest in *Amphiprion*. (Suppl. Info.)

To take into account possible differences in curvature of the vertebral column, we also examined precaudal length by adding the lengths of all precaudal vertebrae following the curvature of the vertebral axis, and divided by sum of the lengths of all vertebrae. Again, we found *Acanthochromis* had the longest precaudal vertebral length of all damselfish genera (p<0.001; Welch One-way ANOVA, Games-Howell post hoc comparisons). *Pomacentrus* was not an exception in this measure, suggesting *Acanthochromis* has a slightly more curved vertebral column that may add additional abdominal space. These measurements show that the consequence of the increase in the number of precaudal vertebrae in *Acanthochromis* is not simply more smaller segments in the same area but an increase in A-P length to the abdominal region. The single additional precaudal vertebra in the other larval brooding genus *Altrichthys* did not result in a detectable difference in abdominal length from other damselfish genera (Suppl. Info. 2). Body cavity depth was variable and not significantly different among taxa (Suppl. Info. 2).

We also measured muscle area (2D area without head, fins, or body cavity). We found that *Acanthochromis* had a significantly smaller muscle area than other genera, and *Lepidozygous* had a significantly larger muscle area, but no significant differences were found among the other genera (Figure 3; Suppl. Info. 2).

### Acanthochromis population differences

Our sample of *Acanthochromis* included specimens from different localities throughout its range. We compared the vertebral numbers and body cavity area from the Philippines, Papua New Guinea, Solomons Islands, Vanuatu, and southern Great Barrier Reef (Suppl. Info.). No differences were detected among populations for muscle area, but southern Great Barrier Reef had a larger body cavity, and Vanuatu had a smaller body cavity (Fig. 2).

### Larval swimming Ucrit comparisons

*Acanthochromis* had a lower Ucrit mean and range than any other damselfish. Once the single *Amphiprion* was removed (i.e. using only species with replicate measurements for statistical tests), the majority of genera are significantly different- and faster than *Acanthochromis*. A one way ANOVA for mean Ucrit between eleven genera of damselfish found that most genera were significantly faster and three genera have a faster mean Ucrit but the difference is not significant (Fig. 4; Suppl Info.2; Fisher and Leis, 2022).

**Figure 4.**
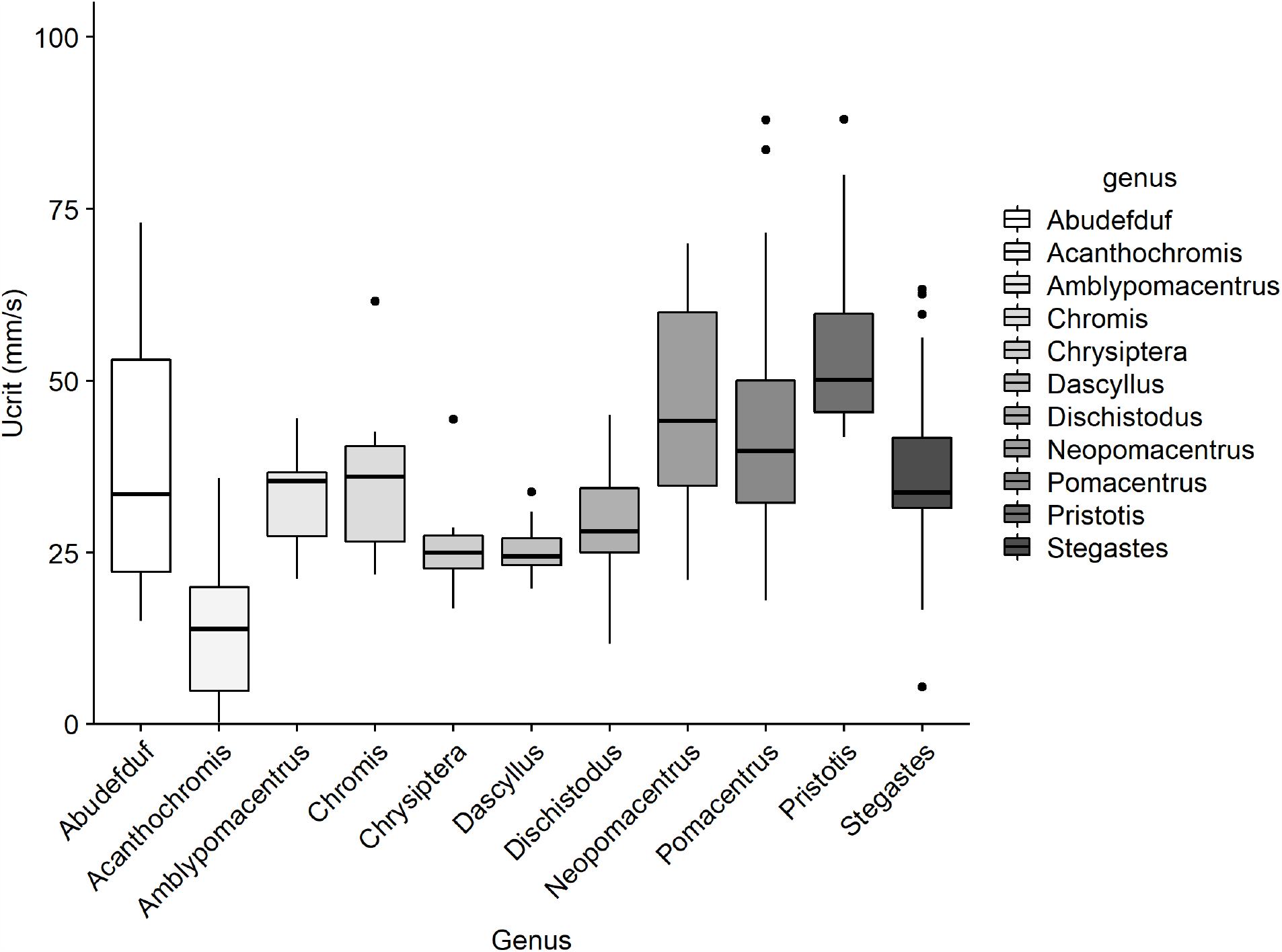
Swimming speeds by pomacentrid genera (taken from the review by Fisher and Leis, 2022). Acanthochromis has the lowest mean Ucrit among pomacentrids.

## Discussion

The consistency in the vertebral count among all damselfishes (except larval brooders) -- including diverse tropical and temperate species, anemonefishes and morphologically divergent damselfish species such as the elongate *Lepidozygous* and the round *Dascyllus-*- suggests a long-stabilized vertebral phenotype in the family. This early stabilization is supported by studies of fossil pomacentrids from the Eocene, which had vertebral counts similar to modern pomacentrids, and no known pomacentrid species has increased numbers of vertebrae beyond XI+15=26 total (Bellwood and Sorbini, 1995). This ∼50 million years of evolution of a diverse speciose family without vertebral count change underlines how extraordinary the precaudal vertebral increase in the larval brooders *Acanthochromis* and *Altrichthys* is. *Acanthochromis polyacanthus* has variably 3-4 additional precaudal vertebrae (Kavanagh, 2000; Leis and Kavanagh, 2003; this study), and here we report for the first time that both *Altrichthys curatus* and *Altrichthys azurelineatus* have one additional precaudal vertebra.

This precaudal expansion in all larval brooding species implies that the lifestyle of this clade is linked to this unusual phenotypic divergence in vertebral count. We previously speculated that the extra precaudal vertebrae indicated selection for a greater body cavity (abdominal) area to hold more of the huge eggs (Kavanagh, 2000). It could be that evolving more precaudal vertebrae could simply result in more, smaller vertebrae but not expand the body cavity. However, here we show the greater number of precaudal vertebrae in *Acanthochromis* (14-15 precaudal vertebrae) is indeed associated with a greater abdominal area. The abdominal area of *Altrichthys s*pp. (12 precaudal vertebrae) is not statistically greater than rest of the typical species with 11 precaudal vertebrae; posthoc analysis found that *Altrichthys* had a bigger abdominal cavity that about half the other genera. The egg size and much of the reproductive biology of *Altrichthys* is unknown, but the reproductive differences between *Altrichthys* and *Acanthochromis* may be informative in understanding functional association with the morphological evolution. One surprising finding was *Pomacentrus* was not statistically different than *Acanthochromis* in abdominal area, although precaudal length measures following the curve of the vertebral column did find differences. Further analysis of this peculiarity in our comparison is necessary for further interpretation.

We tried to discern where the abdominal expansion was happening in the vertebral column by dividing the straight-line abdominal (postcranial precaudal) length into thirds (anterior, middle, posterior) and then counting how many vertebrae were in each third, estimated to the tenth of a vertebra. We found that both *Acanthochromis* and *Altrichthys* have their additional vertebral length distributed roughly equally throughout the precaudal region, suggesting that an overall developmental change in segmentation parameters during early somitogenesis is responsible, rather than a developmental parameter controlling regional growth or segmentation. In contrast, *Lepidozygous*, which is a particularly elongate damselfish, showed a more localized changes in vertebral sizes with larger anterior and smaller posterior precaudal vertebrae, suggesting a different developmental change controlling relative vertebral sizes.

In other fish species, vertebral variation in number and regional boundary shifts are found to vary with temperature, latitude, and sex, and are heritable (Lindsey 1962; Ramler et al., 2014; Yamahira et al., 2006). The nearly invariable pattern of the normal pomacentrid vertebral count over ∼50 million years, despite diversification over a distributional range that extends through tropical, subtropical, and temperate latitudes and habitats, suggests another aspect of their environment must be the stabilizing force.

The pelagic larval stage may hold a clue. The larvae and adults of marine fishes exist in two completely different environments over their life history, described as “two quite separate evolutionary theatres” (Moser and Ahlstrom, 1973). Pelagic larvae have different demands for predator avoidance (Fisher and Leis, 2010), sensory cues (Arvedlund and Kavanagh, 2003), and swimming and feeding challenges (China and Holzman 2014). Sustained distance swimming is more likely to be important adaptation in the pelagic environment for older larvae that need to travel to settlement habitat, while flexibility for turning within complex structural habitats is more likely to be adaptive for adults living on coral reefs (Fisher and Leis, 2010; Schakman and Korsmeyer, 2023). “Pectoral fin swimming” in damselfish adults, rather than tail propulsive swimming, was shown to be among the most efficient energetically for holding position in wave surges on reefs (Schakman and Korsmeyer, 2023).

Laboratory studies of larval reef-fish swimming ability supports the idea that they are capable sustained swimmers. Late stage pelagic fish larvae swim at relatively high speeds (∼0.2-0.6 meters per second or approximately 10-20 body lengths per second; Fisher et al., 2005). Fisher and Bellwood (2002) demonstrated that individual pomacentrid (*Amphiprion melanopus*) larvae can maintain a steady swim for many days and over 16 km nonstop. This late larval swimming is primarily through tail propulsive movement.

Comparisons of swimming in larval marine fishes show that both the amount of tail muscle and the boundary between body cavity (precaudal) and tail (caudal) region affects swimming. The highest contribution to thrust is present in the posterior region, and older larvae use large amplitudes of tail movement to swim (Voesenek et al., 2018). Survival through the pelagic larval stage likely depends on swimming ability.

In larval brooding damselfishes, the “evolutionary theatre” of the pelagic environment is removed, changing the balance of selective forces on many aspects of the phenotype, particularly those determined early in development like vertebral number.

Our observations of muscle area differences between larval brooders and non-brooders further support the idea of a changing selective environment in species without pelagic larvae. We found that *Acanthochromis* had a reduced muscle area, including loss of primarily the posterior muscle used for propulsive tail swimming in pelagic larvae. The muscle area was smaller but the caudal vertebral number was similar to other damselfishes (15).

In a comparison of known larval swimming ability among reef fishes, young juveniles of the larval brooder A*canthochromi*s performed most poorly (Fisher et al., 2022). We propose that when larval brooding evolved, relaxed selection on larval swimming performance allowed a shift in body proportions to favor a larger body cavity and altered axial patterning. We considered that the swimming test may be confounded by the difference in “practice” because *Acanthochromis* spends much longer in the egg and does not have a pelagic stage where thrust swimming is required, however the test specimens were several weeks old, not newly hatched, so those differences were reduced. Enlarged body cavity potentially gives a fitness advantage, as females with a larger abdominal cavity could hold more of the large eggs and increase clutch size. Since *Acanthochromis* invests far more time in parental care of a single brood than other damselfishes, increasing clutch size to increase probability of success makes sense, especially if reducing tail muscle doesn’t compromise swimming on the reef environment.

The lack of a pelagic larval stage also impacts the gene flow among populations, with separated populations showing strong genetic divergence and stable hybrid zones in previous studies (Planes, et al., 2001; Planes and Doherty, 1997). *Acanthochromis* has a distributional range fairly widespread in the Indo-West Pacific. When we examined body proportions by population, we found significant differences in both body cavity area and muscle area among geographically separated populations (Fig. 3). The southern Great Barrier Reef is a genetically isolated population with the coolest seasonal water temperatures (Planes and Doherty, 1997) and that population had both the largest body cavity area and among the smallest muscle area; Vanuatu had the smallest body cavity area (Fig. 3). The fitness consequences of these evolved proportion differences are intriguing and need to be tested for adaptive significance. Both the reproductive biology and the diversification pattern in *Altrichthys* species, a genus endemic to northern Palawan and the Calamianes Islands of the western Philippines, is unknown but could provide further comparative evidence for how lack of a pelagic larval stage changes the selective forces on morphological evolution.

The influence of the pelagic larval stage, and its “separate evolutionary theatre,” on the evolution of the benthic adult morphology is difficult to study even from a comparative view because of the prevalence of pelagic larvae. The damselfishes provide a rare perspective in having closely related species with and without pelagic larvae. This innovation in life history and reproductive biology in the larval brooding damselfishes, and the associated morphological shifts in meristics and body proportions seen in *Acanthochromis*, provides a novel perspective as a rare evolutionary experiment in the marine environment and suggests the pelagic larval environment may constrain morphological evolution of benthic marine fishes.

## Acknowledgements

Specimens and radiographs were kindly provided by the Bernice P. Bishop Museum, Museum of Western Australia, Australian Museum, American Museum of Natural History, and Museum of Comparative Zoology. D. Santiago and L. Pettey provided radiographs. S. Mallick and R. Drew consulted on statistics and graphics. A. Russell and C. Salem helped with morphometrics.

