## Supplemental File 2 for "Larval brooding damselfishes and shifting body proportions: does pelagic larval swimming constrain reef fish morphology?"

A. **Vertebral Counts**

**CV=caudal vertebrae; PCV=precaudal vertebrae**

| **Frequencies for CV Number** | | | | | | | | | | |
| --- | --- | --- | --- | --- | --- | --- | --- | --- | --- | --- |
| **Genus** | **CV Number** | | **Frequency** | | **Percent** | | | **N Percent** | | **Cumulative Percent** |
| Acanthochromis | 14 |  | 6 |  | 4.724 | |  | 4.762 |  | 4.762 |
|  | 15 |  | 120 |  | 94.488 | |  | 95.238 |  | 100 |
|  | Missing |  | 1 |  | 0.787 | |  |  |  |  |
|  | Total |  | 127 |  | 100 | |  |  |  |  |
| Altrichthys | 14 |  | 0 |  | 0 | |  | 0 |  | 0 |
|  | 15 |  | 19 |  | 95 | |  | 100 |  | 100 |
|  | Missing |  | 1 |  | 5 | |  |  |  |  |
|  | Total |  | 20 |  | 100 | |  |  |  |  |
| Amblyglyphidodon | 14 |  | 0 |  | 0 | |  | 0 |  | 0 |
|  | 15 |  | 27 |  | 33.333 | |  | 100 |  | 100 |
|  | Missing |  | 54 |  | 66.667 | |  |  |  |  |
|  | Total |  | 81 |  | 100 | |  |  |  |  |
| Amphiprion | 14 |  | 0 |  | 0 | |  | 0 |  | 0 |
|  | 15 |  | 19 |  | 8.407 | |  | 100 |  | 100 |
|  | Missing |  | 207 |  | 91.593 | |  |  |  |  |
|  | Total |  | 226 |  | 100 | |  |  |  |  |
| Chromis | 14 |  | 0 |  | 0 | |  | 0 |  | 0 |
|  | 15 |  | 16 |  | 88.889 | |  | 100 |  | 100 |
|  | Missing |  | 2 |  | 11.111 | |  |  |  |  |
|  | Total |  | 18 |  | 100 | |  |  |  |  |
| Dascyllus | 14 |  | 0 |  | 0 | |  | 0 |  | 0 |
|  | 15 |  | 17 |  | 100 | |  | 100 |  | 100 |
|  | Missing |  | 0 |  | 0 | |  |  |  |  |
|  | Total |  | 17 |  | 100 | |  |  |  |  |
| Hemiglyphidodon | 14 |  | 0 |  | 0 | |  | 0 |  | 0 |
|  | 15 |  | 37 |  | 88.095 | |  | 100 |  | 100 |
|  | Missing |  | 5 |  | 11.905 | |  |  |  |  |
|  | Total |  | 42 |  | 100 | |  |  |  |  |
| Lepidozygus | 14 |  | 0 |  | 0 | |  | 0 |  | 0 |
|  | 15 |  | 16 |  | 100 | |  | 100 |  | 100 |
|  | Missing |  | 0 |  | 0 | |  |  |  |  |
|  | Total |  | 16 |  | 100 | |  |  |  |  |
| Neoglyphidodon | 14 |  | 0 |  | 0 | |  | 0 |  | 0 |
|  | 15 |  | 26 |  | 38.806 | |  | 100 |  | 100 |
|  | Missing |  | 41 |  | 61.194 | |  |  |  |  |
|  | Total |  | 67 |  | 100 | |  |  |  |  |
| Pomacentrus | 14 |  | 0 |  | 0 | |  | 0 |  | 0 |
|  | 15 |  | 5 |  | 100 | |  | 100 |  | 100 |
|  | Missing |  | 0 |  | 0 | |  |  |  |  |
|  | Total |  | 5 |  | 100 | |  |  |  |  |
| **Frequencies for PCV Number** | | | | | | | | | | |
| **Genus** | **PCV Number** | | **Frequency** | | **Percent** | | | **N Percent** | | **Cumulative Percent** |
| Acanthochromis | 10 |  | 0 | |  | 0 |  | 0 |  | 0 |
|  | 11 |  | 0 | |  | 0 |  | 0 |  | 0 |
|  | 12 |  | 0 | |  | 0 |  | 0 |  | 0 |
|  | 13 |  | 2 | |  | 1.575 |  | 1.587 |  | 1.587 |
|  | 14 |  | 87 | |  | 68.504 |  | 69.048 |  | 70.635 |
|  | 15 |  | 37 | |  | 29.134 |  | 29.365 |  | 100 |
|  | Missing |  | 1 | |  | 0.787 |  |  |  |  |
|  | Total |  | 127 | |  | 100 |  |  |  |  |
| Altrichthys | 10 |  | 0 | |  | 0 |  | 0 |  | 0 |
|  | 11 |  | 1 | |  | 5 |  | 5.263 |  | 5.263 |
|  | 12 |  | 18 | |  | 90 |  | 94.737 |  | 100 |
|  | 13 |  | 0 | |  | 0 |  | 0 |  | 100 |
|  | 14 |  | 0 | |  | 0 |  | 0 |  | 100 |
|  | 15 |  | 0 | |  | 0 |  | 0 |  | 100 |
|  | Missing |  | 1 | |  | 5 |  |  |  |  |
|  | Total |  | 20 | |  | 100 |  |  |  |  |
| Amblyglyphidodon | 10 |  | 1 | |  | 1.235 |  | 3.704 |  | 3.704 |
|  | 11 |  | 26 | |  | 32.099 |  | 96.296 |  | 100 |
|  | 12 |  | 0 | |  | 0 |  | 0 |  | 100 |
|  | 13 |  | 0 | |  | 0 |  | 0 |  | 100 |
|  | 14 |  | 0 | |  | 0 |  | 0 |  | 100 |
|  | 15 |  | 0 | |  | 0 |  | 0 |  | 100 |
|  | Missing |  | 54 | |  | 66.667 |  |  |  |  |
|  | Total |  | 81 | |  | 100 |  |  |  |  |
| Amphiprion | 10 |  | 0 | |  | 0 |  | 0 |  | 0 |
|  | 11 |  | 19 | |  | 8.407 |  | 100 |  | 100 |
|  | 12 |  | 0 | |  | 0 |  | 0 |  | 100 |
|  | 13 |  | 0 | |  | 0 |  | 0 |  | 100 |
|  | 14 |  | 0 | |  | 0 |  | 0 |  | 100 |
|  | 15 |  | 0 | |  | 0 |  | 0 |  | 100 |
|  | Missing |  | 207 | |  | 91.593 |  |  |  |  |
|  | Total |  | 226 | |  | 100 |  |  |  |  |
| Chromis | 10 |  | 0 | |  | 0 |  | 0 |  | 0 |
|  | 11 |  | 16 | |  | 88.889 |  | 100 |  | 100 |
|  | 12 |  | 0 | |  | 0 |  | 0 |  | 100 |
|  | 13 |  | 0 | |  | 0 |  | 0 |  | 100 |
|  | 14 |  | 0 | |  | 0 |  | 0 |  | 100 |
|  | 15 |  | 0 | |  | 0 |  | 0 |  | 100 |
|  | Missing |  | 2 | |  | 11.111 |  |  |  |  |
|  | Total |  | 18 | |  | 100 |  |  |  |  |
| Dascyllus | 10 |  | 0 | |  | 0 |  | 0 |  | 0 |
|  | 11 |  | 17 | |  | 100 |  | 100 |  | 100 |
|  | 12 |  | 0 | |  | 0 |  | 0 |  | 100 |
|  | 13 |  | 0 | |  | 0 |  | 0 |  | 100 |
|  | 14 |  | 0 | |  | 0 |  | 0 |  | 100 |
|  | 15 |  | 0 | |  | 0 |  | 0 |  | 100 |
|  | Missing |  | 0 | |  | 0 |  |  |  |  |
|  | Total |  | 17 | |  | 100 |  |  |  |  |
| Hemiglyphidodon | 10 |  | 0 | |  | 0 |  | 0 |  | 0 |
|  | 11 |  | 37 | |  | 88.095 |  | 100 |  | 100 |
|  | 12 |  | 0 | |  | 0 |  | 0 |  | 100 |
|  | 13 |  | 0 | |  | 0 |  | 0 |  | 100 |
|  | 14 |  | 0 | |  | 0 |  | 0 |  | 100 |
|  | 15 |  | 0 | |  | 0 |  | 0 |  | 100 |
|  | Missing |  | 5 | |  | 11.905 |  |  |  |  |
|  | Total |  | 42 | |  | 100 |  |  |  |  |
| Lepidozygus | 10 |  | 1 | |  | 6.25 |  | 6.25 |  | 6.25 |
|  | 11 |  | 15 | |  | 93.75 |  | 93.75 |  | 100 |
|  | 12 |  | 0 | |  | 0 |  | 0 |  | 100 |
|  | 13 |  | 0 | |  | 0 |  | 0 |  | 100 |
|  | 14 |  | 0 | |  | 0 |  | 0 |  | 100 |
|  | 15 |  | 0 | |  | 0 |  | 0 |  | 100 |
|  | Missing |  | 0 | |  | 0 |  |  |  |  |
|  | Total |  | 16 | |  | 100 |  |  |  |  |
| Neoglyphidodon | 10 |  | 0 | |  | 0 |  | 0 |  | 0 |
|  | 11 |  | 26 | |  | 38.806 |  | 100 |  | 100 |
|  | 12 |  | 0 | |  | 0 |  | 0 |  | 100 |
|  | 13 |  | 0 | |  | 0 |  | 0 |  | 100 |
|  | 14 |  | 0 | |  | 0 |  | 0 |  | 100 |
|  | 15 |  | 0 | |  | 0 |  | 0 |  | 100 |
|  | Missing |  | 41 | |  | 61.194 |  |  |  |  |
|  | Total |  | 67 | |  | 100 |  |  |  |  |
| Pomacentrus | 10 |  | 0 | |  | 0 |  | 0 |  | 0 |
|  | 11 |  | 5 | |  | 100 |  | 100 |  | 100 |
|  | 12 |  | 0 | |  | 0 |  | 0 |  | 100 |
|  | 13 |  | 0 | |  | 0 |  | 0 |  | 100 |
|  | 14 |  | 0 | |  | 0 |  | 0 |  | 100 |
|  | 15 |  | 0 | |  | 0 |  | 0 |  | 100 |
|  | Missing |  | 0 | |  | 0 |  |  |  |  |
|  | Total |  | 5 | |  | 100 |  |  |  |  |

**Precaudal vertebral “thirds” counts**

Means and ranges of number of vertebrae in each third of the precaudal region, measured on straight-line A-P axis. Number of vertebrae estimated to the nearest tenth of a vertebra within the space.

| Taxon | Anterior | Middle | Posterior | Vert. Formula |
| --- | --- | --- | --- | --- |
| *Acanthochromis polyacanthus** | 6.18 (5.9 – 6.4) | 4.38 (4.2 – 4.5) | 3.98 (3.8 – 4.1) | XV+15=30^#^ |
| *Altrichthys curatus** | 5.18 (5.1 - 5.3) | 3.54 (3.5 – 3.6) | 3.32 (3.1 – 3.8) | XII+15=27 |
| *Amblyglyphididon curacao* | 4.82 (4.6 – 5.0) | 3.18 (3.1 – 3.3) | 3.02 (3.0 – 3.1) | XI+15=26 |
| *Dascyllus trimaculatus* | 4.90 (4.6 – 5.2) | 3.12 (2.9 – 3.4) | 2.98 (2.9 – 3.0) | XI+15=26 |
| *Lepidozygous tapeinosoma* | 4.46 (4.2 – 4.7) | 3.20 (2.4 – 3.6) | 3.78 (3.2 – 4.9) | XI+15=26 |

**Acanthochromis population vertebral count frequency**

PCV=Precaudal vertebrae; CV=Caudal vertebrae; TotV=Total vertebrae

| Location | N | **PCV#** | Freq | **CV#** | Freq | **TotV#** | Freq |
| --- | --- | --- | --- | --- | --- | --- | --- |
| Philippines | 5 | **14** | 0.8 | **15** | 1.0 | **29** | 0.8 |
|  |  | **15** | 0.2 |  |  | **30** | 0.2 |
| Papua NG | 28 | **15** | 1.0 | **13** | 0.07 (2/28) | **28** | 0.07 |
|  |  |  |  | **14** | 0.68 (19/28) | **29** | 0.68 |
|  |  |  |  | **15** | 0.25 (7/28) | **30** | 0.25 |
| Solomon | 23 | **15** | 1.0 | **14** | 0.48 (11/23) | **29** | 0.48 |
|  |  |  |  | **15** | 0.52 (12/23) | **30** | 0.52 |
| Vanuatu | 19 | **15** | 1.0 | **14** | 0.95 | **29** | 0.95 |
|  |  | **15** | 0.05 | **30** | 0.05 |  |  |
| S-GBR | 49 | **14** | 0.04 | **14** | 0.76 (37/49) | **28** | 0.04 |
|  |  | **15** | 0.96 | **15** | 0.24 (12/49) | **29** | 0.71 |
|  |  |  |  |  |  | **30** | 0.24 |

B. **Body area proportions**

ANOVA for Body Cavity Area / Total Body Area:

| **ANOVA - BCA/TBA** | | | | | | | | | | | | |
| --- | --- | --- | --- | --- | --- | --- | --- | --- | --- | --- | --- | --- |
| **Cases** | | **Homogeneity Correction** | | **Sum of Squares** | | **df** | | **Mean Square** | | **F** | | **p** |
| Genus |  | Welch |  | 0.341 |  | 9.000 |  | 0.038 |  | 102.631 |  | < .001 |
| Residual |  | Welch |  | 0.205 |  | 64.973 |  | 0.003 |  |  |  |  |
| *Note.*  Type III Sum of Squares | | | | | | | | | | | | |

| **Test for Equality of Variances (Levene's)** | | | | | | |
| --- | --- | --- | --- | --- | --- | --- |
| **F** | | **df1** | | **df2** | | **p** |
| 5.923 |  | 9.000 |  | 609.000 |  | < .001 |

**Post Hoc Tests**

| **Games-Howell Post Hoc Comparisons - Genus** | | | | | | | | | | |
| --- | --- | --- | --- | --- | --- | --- | --- | --- | --- | --- |
|  | |  | | **Mean Difference** | | **SE** | | **t** | | **p** |
| Acanthochromis |  | Altrichthys |  | 0.025 |  | 0.003 |  | 9.919 |  | < .001 |
|  |  | Amblyglyphidodon |  | 0.036 |  | 0.003 |  | 14.247 |  | < .001 |
|  |  | Amphiprion |  | 0.059 |  | 0.002 |  | 26.201 |  | < .001 |
|  |  | Chromis |  | 0.040 |  | 0.004 |  | 9.208 |  | < .001 |
|  |  | Dascyllus |  | 0.033 |  | 0.005 |  | 6.601 |  | < .001 |
|  |  | Hemiglyphidodon |  | 0.023 |  | 0.003 |  | 8.401 |  | < .001 |
|  |  | Lepidozygus |  | 0.072 |  | 0.004 |  | 18.003 |  | < .001 |
|  |  | Neoglyphidodon |  | 0.018 |  | 0.004 |  | 5.044 |  | < .001 |
|  |  | Pomacentrus |  | 0.017 |  | 0.006 |  | 3.079 |  | 0.248 |
| Altrichthys |  | Amblyglyphidodon |  | 0.011 |  | 0.002 |  | 4.692 |  | < .001 |
|  |  | Amphiprion |  | 0.034 |  | 0.002 |  | 17.065 |  | < .001 |
|  |  | Chromis |  | 0.015 |  | 0.004 |  | 3.542 |  | 0.045 |
|  |  | Dascyllus |  | 0.007 |  | 0.005 |  | 1.541 |  | 0.860 |
|  |  | Hemiglyphidodon |  | -0.002 |  | 0.002 |  | -0.981 |  | 0.992 |
|  |  | Lepidozygus |  | 0.047 |  | 0.004 |  | 12.164 |  | < .001 |
|  |  | Neoglyphidodon |  | -0.007 |  | 0.003 |  | -2.148 |  | 0.500 |
|  |  | Pomacentrus |  | -0.008 |  | 0.006 |  | -1.410 |  | 0.881 |
| Amblyglyphidodon |  | Amphiprion |  | 0.023 |  | 0.002 |  | 11.837 |  | < .001 |
|  |  | Chromis |  | 0.004 |  | 0.004 |  | 1.009 |  | 0.989 |
|  |  | Dascyllus |  | -0.003 |  | 0.005 |  | -0.684 |  | 0.999 |
|  |  | Hemiglyphidodon |  | -0.013 |  | 0.002 |  | -5.329 |  | < .001 |
|  |  | Lepidozygus |  | 0.036 |  | 0.004 |  | 9.423 |  | < .001 |
|  |  | Neoglyphidodon |  | -0.018 |  | 0.003 |  | -5.337 |  | < .001 |
|  |  | Pomacentrus |  | -0.019 |  | 0.006 |  | -3.351 |  | 0.204 |
| Amphiprion |  | Chromis |  | -0.019 |  | 0.004 |  | -4.701 |  | 0.004 |
|  |  | Dascyllus |  | -0.027 |  | 0.005 |  | -5.696 |  | < .001 |
|  |  | Hemiglyphidodon |  | -0.037 |  | 0.002 |  | -16.554 |  | < .001 |
|  |  | Lepidozygus |  | 0.013 |  | 0.004 |  | 3.510 |  | 0.057 |
|  |  | Neoglyphidodon |  | -0.041 |  | 0.003 |  | -12.996 |  | < .001 |
|  |  | Pomacentrus |  | -0.042 |  | 0.005 |  | -7.739 |  | 0.011 |
| Chromis |  | Dascyllus |  | -0.008 |  | 0.006 |  | -1.261 |  | 0.955 |
|  |  | Hemiglyphidodon |  | -0.017 |  | 0.004 |  | -4.022 |  | 0.014 |
|  |  | Lepidozygus |  | 0.032 |  | 0.005 |  | 6.117 |  | < .001 |
|  |  | Neoglyphidodon |  | -0.022 |  | 0.005 |  | -4.539 |  | 0.002 |
|  |  | Pomacentrus |  | -0.023 |  | 0.007 |  | -3.465 |  | 0.111 |
| Dascyllus |  | Hemiglyphidodon |  | -0.010 |  | 0.005 |  | -2.009 |  | 0.602 |
|  |  | Lepidozygus |  | 0.040 |  | 0.006 |  | 6.911 |  | < .001 |
|  |  | Neoglyphidodon |  | -0.015 |  | 0.005 |  | -2.711 |  | 0.213 |
|  |  | Pomacentrus |  | -0.015 |  | 0.007 |  | -2.185 |  | 0.515 |
| Hemiglyphidodon |  | Lepidozygus |  | 0.050 |  | 0.004 |  | 12.435 |  | < .001 |
|  |  | Neoglyphidodon |  | -0.005 |  | 0.004 |  | -1.375 |  | 0.932 |
|  |  | Pomacentrus |  | -0.005 |  | 0.006 |  | -0.955 |  | 0.984 |
| Lepidozygus |  | Neoglyphidodon |  | -0.054 |  | 0.005 |  | -11.826 |  | < .001 |
|  |  | Pomacentrus |  | -0.055 |  | 0.006 |  | -8.644 |  | < .001 |
| Neoglyphidodon |  | Pomacentrus |  | -5.330e -4 |  | 0.006 |  | -0.088 |  | 1.000 |

BCA / (Total Body Area – Head):

| **ANOVA - Cavity / (BodyArea - Head)** | | | | | | | | | | | | |
| --- | --- | --- | --- | --- | --- | --- | --- | --- | --- | --- | --- | --- |
| **Cases** | | **Homogeneity Correction** | | **Sum of Squares** | | **df** | | **Mean Square** | | **F** | | **p** |
| Genus |  | Welch |  | 0.354 |  | 9.000 |  | 0.039 |  | 73.145 |  | < .001 |
| Residual |  | Welch |  | 0.303 |  | 64.950 |  | 0.005 |  |  |  |  |
| *Note.*  Type III Sum of Squares | | | | | | | | | | | | |

| **Test for Equality of Variances (Levene's)** | | | | | | |
| --- | --- | --- | --- | --- | --- | --- |
| **F** | | **df1** | | **df2** | | **p** |
| 5.966 |  | 9.000 |  | 609.000 |  | < .001 |

| **Games-Howell Post Hoc Comparisons - Genus** | | | | | | | | | | |
| --- | --- | --- | --- | --- | --- | --- | --- | --- | --- | --- |
|  | |  | | **Mean Difference** | | **SE** | | **t** | | **p** |
| Acanthochromis |  | Altrichthys |  | 0.031 |  | 0.003 |  | 9.028 |  | < .001 |
|  |  | Amblyglyphidodon |  | 0.046 |  | 0.003 |  | 14.710 |  | < .001 |
|  |  | Amphiprion |  | 0.057 |  | 0.003 |  | 20.187 |  | < .001 |
|  |  | Chromis |  | 0.050 |  | 0.005 |  | 9.306 |  | < .001 |
|  |  | Dascyllus |  | 0.036 |  | 0.005 |  | 6.609 |  | < .001 |
|  |  | Hemiglyphidodon |  | 0.016 |  | 0.004 |  | 4.419 |  | < .001 |
|  |  | Lepidozygus |  | 0.086 |  | 0.005 |  | 18.928 |  | < .001 |
|  |  | Neoglyphidodon |  | 0.018 |  | 0.004 |  | 4.399 |  | < .001 |
|  |  | Pomacentrus |  | 0.026 |  | 0.006 |  | 4.067 |  | 0.092 |
| Altrichthys |  | Amblyglyphidodon |  | 0.014 |  | 0.003 |  | 4.551 |  | 0.002 |
|  |  | Amphiprion |  | 0.026 |  | 0.003 |  | 9.047 |  | < .001 |
|  |  | Chromis |  | 0.019 |  | 0.005 |  | 3.487 |  | 0.047 |
|  |  | Dascyllus |  | 0.005 |  | 0.005 |  | 0.882 |  | 0.996 |
|  |  | Hemiglyphidodon |  | -0.015 |  | 0.004 |  | -4.251 |  | 0.003 |
|  |  | Lepidozygus |  | 0.055 |  | 0.005 |  | 12.013 |  | < .001 |
|  |  | Neoglyphidodon |  | -0.013 |  | 0.004 |  | -3.080 |  | 0.079 |
|  |  | Pomacentrus |  | -0.006 |  | 0.006 |  | -0.923 |  | 0.987 |
| Amblyglyphidodon |  | Amphiprion |  | 0.012 |  | 0.002 |  | 4.907 |  | < .001 |
|  |  | Chromis |  | 0.005 |  | 0.005 |  | 0.914 |  | 0.994 |
|  |  | Dascyllus |  | -0.009 |  | 0.005 |  | -1.767 |  | 0.747 |
|  |  | Hemiglyphidodon |  | -0.030 |  | 0.003 |  | -9.062 |  | < .001 |
|  |  | Lepidozygus |  | 0.041 |  | 0.004 |  | 9.509 |  | < .001 |
|  |  | Neoglyphidodon |  | -0.027 |  | 0.004 |  | -6.956 |  | < .001 |
|  |  | Pomacentrus |  | -0.020 |  | 0.006 |  | -3.272 |  | 0.215 |
| Amphiprion |  | Chromis |  | -0.007 |  | 0.005 |  | -1.358 |  | 0.926 |
|  |  | Dascyllus |  | -0.021 |  | 0.005 |  | -4.080 |  | 0.018 |
|  |  | Hemiglyphidodon |  | -0.041 |  | 0.003 |  | -13.663 |  | < .001 |
|  |  | Lepidozygus |  | 0.029 |  | 0.004 |  | 7.129 |  | < .001 |
|  |  | Neoglyphidodon |  | -0.039 |  | 0.004 |  | -10.484 |  | < .001 |
|  |  | Pomacentrus |  | -0.031 |  | 0.006 |  | -5.278 |  | 0.048 |
| Chromis |  | Dascyllus |  | -0.014 |  | 0.007 |  | -2.039 |  | 0.580 |
|  |  | Hemiglyphidodon |  | -0.034 |  | 0.005 |  | -6.234 |  | < .001 |
|  |  | Lepidozygus |  | 0.036 |  | 0.006 |  | 5.836 |  | < .001 |
|  |  | Neoglyphidodon |  | -0.032 |  | 0.006 |  | -5.395 |  | < .001 |
|  |  | Pomacentrus |  | -0.025 |  | 0.008 |  | -3.275 |  | 0.130 |
| Dascyllus |  | Hemiglyphidodon |  | -0.020 |  | 0.006 |  | -3.630 |  | 0.033 |
|  |  | Lepidozygus |  | 0.050 |  | 0.006 |  | 8.017 |  | < .001 |
|  |  | Neoglyphidodon |  | -0.018 |  | 0.006 |  | -2.979 |  | 0.124 |
|  |  | Pomacentrus |  | -0.011 |  | 0.008 |  | -1.403 |  | 0.902 |
| Hemiglyphidodon |  | Lepidozygus |  | 0.070 |  | 0.005 |  | 15.040 |  | < .001 |
|  |  | Neoglyphidodon |  | 0.002 |  | 0.004 |  | 0.571 |  | 1.000 |
|  |  | Pomacentrus |  | 0.010 |  | 0.006 |  | 1.506 |  | 0.851 |
| Lepidozygus |  | Neoglyphidodon |  | -0.068 |  | 0.005 |  | -13.204 |  | < .001 |
|  |  | Pomacentrus |  | -0.061 |  | 0.007 |  | -8.716 |  | < .001 |
| Neoglyphidodon |  | Pomacentrus |  | 0.007 |  | 0.007 |  | 1.061 |  | 0.975 |

Muscle Area / (Total Body Area – Head):

| **ANOVA - Muscle Area / (BodyArea - Head)** | | | | | | | | | | | | |
| --- | --- | --- | --- | --- | --- | --- | --- | --- | --- | --- | --- | --- |
| **Cases** | | **Homogeneity Correction** | | **Sum of Squares** | | **df** | | **Mean Square** | | **F** | | **p** |
| Genus |  | Welch |  | 0.354 |  | 9.000 |  | 0.039 |  | 73.145 |  | < .001 |
| Residual |  | Welch |  | 0.303 |  | 64.950 |  | 0.005 |  |  |  |  |
| *Note.*  Type III Sum of Squares | | | | | | | | | | | | |

| **Test for Equality of Variances (Levene's)** | | | | | | |
| --- | --- | --- | --- | --- | --- | --- |
| **F** | | **df1** | | **df2** | | **p** |
| 5.966 |  | 9.000 |  | 609.000 |  | < .001 |

**Post Hoc Tests**

| **Games-Howell Post Hoc Comparisons - Genus** | | | | | | | | | | |
| --- | --- | --- | --- | --- | --- | --- | --- | --- | --- | --- |
|  | |  | | **Mean Difference** | | **SE** | | **t** | | **p** |
| Acanthochromis |  | Altrichthys |  | -0.031 |  | 0.003 |  | -9.028 |  | < .001 |
|  |  | Amblyglyphidodon |  | -0.046 |  | 0.003 |  | -14.710 |  | < .001 |
|  |  | Amphiprion |  | -0.057 |  | 0.003 |  | -20.187 |  | < .001 |
|  |  | Chromis |  | -0.050 |  | 0.005 |  | -9.306 |  | < .001 |
|  |  | Dascyllus |  | -0.036 |  | 0.005 |  | -6.609 |  | < .001 |
|  |  | Hemiglyphidodon |  | -0.016 |  | 0.004 |  | -4.419 |  | < .001 |
|  |  | Lepidozygus |  | -0.086 |  | 0.005 |  | -18.928 |  | < .001 |
|  |  | Neoglyphidodon |  | -0.018 |  | 0.004 |  | -4.399 |  | < .001 |
|  |  | Pomacentrus |  | -0.026 |  | 0.006 |  | -4.067 |  | 0.092 |
| Altrichthys |  | Amblyglyphidodon |  | -0.014 |  | 0.003 |  | -4.551 |  | 0.002 |
|  |  | Amphiprion |  | -0.026 |  | 0.003 |  | -9.047 |  | < .001 |
|  |  | Chromis |  | -0.019 |  | 0.005 |  | -3.487 |  | 0.047 |
|  |  | Dascyllus |  | -0.005 |  | 0.005 |  | -0.882 |  | 0.996 |
|  |  | Hemiglyphidodon |  | 0.015 |  | 0.004 |  | 4.251 |  | 0.003 |
|  |  | Lepidozygus |  | -0.055 |  | 0.005 |  | -12.013 |  | < .001 |
|  |  | Neoglyphidodon |  | 0.013 |  | 0.004 |  | 3.080 |  | 0.079 |
|  |  | Pomacentrus |  | 0.006 |  | 0.006 |  | 0.923 |  | 0.987 |
| Amblyglyphidodon |  | Amphiprion |  | -0.012 |  | 0.002 |  | -4.907 |  | < .001 |
|  |  | Chromis |  | -0.005 |  | 0.005 |  | -0.914 |  | 0.994 |
|  |  | Dascyllus |  | 0.009 |  | 0.005 |  | 1.767 |  | 0.747 |
|  |  | Hemiglyphidodon |  | 0.030 |  | 0.003 |  | 9.062 |  | < .001 |
|  |  | Lepidozygus |  | -0.041 |  | 0.004 |  | -9.509 |  | < .001 |
|  |  | Neoglyphidodon |  | 0.027 |  | 0.004 |  | 6.956 |  | < .001 |
|  |  | Pomacentrus |  | 0.020 |  | 0.006 |  | 3.272 |  | 0.215 |
| Amphiprion |  | Chromis |  | 0.007 |  | 0.005 |  | 1.358 |  | 0.926 |
|  |  | Dascyllus |  | 0.021 |  | 0.005 |  | 4.080 |  | 0.018 |
|  |  | Hemiglyphidodon |  | 0.041 |  | 0.003 |  | 13.663 |  | < .001 |
|  |  | Lepidozygus |  | -0.029 |  | 0.004 |  | -7.129 |  | < .001 |
|  |  | Neoglyphidodon |  | 0.039 |  | 0.004 |  | 10.484 |  | < .001 |
|  |  | Pomacentrus |  | 0.031 |  | 0.006 |  | 5.278 |  | 0.048 |
| Chromis |  | Dascyllus |  | 0.014 |  | 0.007 |  | 2.039 |  | 0.580 |
|  |  | Hemiglyphidodon |  | 0.034 |  | 0.005 |  | 6.234 |  | < .001 |
|  |  | Lepidozygus |  | -0.036 |  | 0.006 |  | -5.836 |  | < .001 |
|  |  | Neoglyphidodon |  | 0.032 |  | 0.006 |  | 5.395 |  | < .001 |
|  |  | Pomacentrus |  | 0.025 |  | 0.008 |  | 3.275 |  | 0.130 |
| Dascyllus |  | Hemiglyphidodon |  | 0.020 |  | 0.006 |  | 3.630 |  | 0.033 |
|  |  | Lepidozygus |  | -0.050 |  | 0.006 |  | -8.017 |  | < .001 |
|  |  | Neoglyphidodon |  | 0.018 |  | 0.006 |  | 2.979 |  | 0.124 |
|  |  | Pomacentrus |  | 0.011 |  | 0.008 |  | 1.403 |  | 0.902 |
| Hemiglyphidodon |  | Lepidozygus |  | -0.070 |  | 0.005 |  | -15.040 |  | < .001 |
|  |  | Neoglyphidodon |  | -0.002 |  | 0.004 |  | -0.571 |  | 1.000 |
|  |  | Pomacentrus |  | -0.010 |  | 0.006 |  | -1.506 |  | 0.851 |
| Lepidozygus |  | Neoglyphidodon |  | 0.068 |  | 0.005 |  | 13.204 |  | < .001 |
|  |  | Pomacentrus |  | 0.061 |  | 0.007 |  | 8.716 |  | < .001 |
| Neoglyphidodon |  | Pomacentrus |  | -0.007 |  | 0.007 |  | -1.061 |  | 0.975 |

Head Area / Total Body Area:

| **ANOVA - Head/TBA** | | | | | | | | | | | | |
| --- | --- | --- | --- | --- | --- | --- | --- | --- | --- | --- | --- | --- |
| **Cases** | | **Homogeneity Correction** | | **Sum of Squares** | | **df** | | **Mean Square** | | **F** | | **p** |
| Genus |  | Welch |  | 0.330 |  | 9.000 |  | 0.037 |  | 76.458 |  | < .001 |
| Residual |  | Welch |  | 0.288 |  | 67.981 |  | 0.004 |  |  |  |  |
| *Note.*  Type III Sum of Squares | | | | | | | | | | | | |

| **Test for Equality of Variances (Levene's)** | | | | | | |
| --- | --- | --- | --- | --- | --- | --- |
| **F** | | **df1** | | **df2** | | **p** |
| 9.958 |  | 9.000 |  | 609.000 |  | < .001 |

| **Games-Howell Post Hoc Comparisons - Genus** | | | | | | | | | | |
| --- | --- | --- | --- | --- | --- | --- | --- | --- | --- | --- |
|  | |  | | **Mean Difference** | | **SE** | | **t** | | **p** |
| Acanthochromis |  | Altrichthys |  | -0.003 |  | 0.002 |  | -1.263 |  | 0.957 |
|  |  | Amblyglyphidodon |  | -0.003 |  | 0.003 |  | -1.114 |  | 0.983 |
|  |  | Amphiprion |  | -0.053 |  | 0.002 |  | -22.883 |  | < .001 |
|  |  | Chromis |  | -0.005 |  | 0.003 |  | -2.026 |  | 0.587 |
|  |  | Dascyllus |  | -0.016 |  | 0.005 |  | -3.055 |  | 0.131 |
|  |  | Hemiglyphidodon |  | -0.031 |  | 0.003 |  | -10.174 |  | < .001 |
|  |  | Lepidozygus |  | -0.023 |  | 0.007 |  | -3.301 |  | 0.091 |
|  |  | Neoglyphidodon |  | -0.012 |  | 0.003 |  | -3.758 |  | 0.010 |
|  |  | Pomacentrus |  | 0.007 |  | 0.003 |  | 2.014 |  | 0.619 |
| Altrichthys |  | Amblyglyphidodon |  | 1.490e -4 |  | 0.003 |  | 0.053 |  | 1.000 |
|  |  | Amphiprion |  | -0.050 |  | 0.003 |  | -19.308 |  | < .001 |
|  |  | Chromis |  | -0.002 |  | 0.003 |  | -0.752 |  | 0.999 |
|  |  | Dascyllus |  | -0.013 |  | 0.005 |  | -2.399 |  | 0.374 |
|  |  | Hemiglyphidodon |  | -0.028 |  | 0.003 |  | -8.594 |  | < .001 |
|  |  | Lepidozygus |  | -0.020 |  | 0.007 |  | -2.823 |  | 0.202 |
|  |  | Neoglyphidodon |  | -0.009 |  | 0.003 |  | -2.611 |  | 0.230 |
|  |  | Pomacentrus |  | 0.010 |  | 0.004 |  | 2.741 |  | 0.303 |
| Amblyglyphidodon |  | Amphiprion |  | -0.050 |  | 0.003 |  | -18.120 |  | < .001 |
|  |  | Chromis |  | -0.002 |  | 0.003 |  | -0.760 |  | 0.999 |
|  |  | Dascyllus |  | -0.013 |  | 0.005 |  | -2.386 |  | 0.378 |
|  |  | Hemiglyphidodon |  | -0.028 |  | 0.003 |  | -8.280 |  | < .001 |
|  |  | Lepidozygus |  | -0.020 |  | 0.007 |  | -2.817 |  | 0.202 |
|  |  | Neoglyphidodon |  | -0.009 |  | 0.003 |  | -2.549 |  | 0.253 |
|  |  | Pomacentrus |  | 0.010 |  | 0.004 |  | 2.610 |  | 0.335 |
| Amphiprion |  | Chromis |  | 0.048 |  | 0.003 |  | 17.690 |  | < .001 |
|  |  | Dascyllus |  | 0.037 |  | 0.005 |  | 7.192 |  | < .001 |
|  |  | Hemiglyphidodon |  | 0.022 |  | 0.003 |  | 6.801 |  | < .001 |
|  |  | Lepidozygus |  | 0.030 |  | 0.007 |  | 4.298 |  | 0.013 |
|  |  | Neoglyphidodon |  | 0.041 |  | 0.003 |  | 12.647 |  | < .001 |
|  |  | Pomacentrus |  | 0.060 |  | 0.004 |  | 16.603 |  | < .001 |
| Chromis |  | Dascyllus |  | -0.010 |  | 0.005 |  | -1.977 |  | 0.622 |
|  |  | Hemiglyphidodon |  | -0.026 |  | 0.003 |  | -7.732 |  | < .001 |
|  |  | Lepidozygus |  | -0.018 |  | 0.007 |  | -2.509 |  | 0.326 |
|  |  | Neoglyphidodon |  | -0.007 |  | 0.003 |  | -1.928 |  | 0.650 |
|  |  | Pomacentrus |  | 0.012 |  | 0.004 |  | 3.237 |  | 0.165 |
| Dascyllus |  | Hemiglyphidodon |  | -0.016 |  | 0.006 |  | -2.796 |  | 0.189 |
|  |  | Lepidozygus |  | -0.007 |  | 0.008 |  | -0.872 |  | 0.996 |
|  |  | Neoglyphidodon |  | 0.004 |  | 0.006 |  | 0.691 |  | 0.999 |
|  |  | Pomacentrus |  | 0.023 |  | 0.006 |  | 3.894 |  | 0.026 |
| Hemiglyphidodon |  | Lepidozygus |  | 0.008 |  | 0.007 |  | 1.139 |  | 0.974 |
|  |  | Neoglyphidodon |  | 0.019 |  | 0.004 |  | 5.077 |  | < .001 |
|  |  | Pomacentrus |  | 0.038 |  | 0.004 |  | 9.242 |  | < .001 |
| Lepidozygus |  | Neoglyphidodon |  | 0.011 |  | 0.007 |  | 1.527 |  | 0.866 |
|  |  | Pomacentrus |  | 0.030 |  | 0.007 |  | 4.006 |  | 0.020 |
| Neoglyphidodon |  | Pomacentrus |  | 0.019 |  | 0.004 |  | 4.492 |  | 0.017 |

Body Cavity Length / Standard Length:

| **ANOVA - Body Cavity Length / SL** | | | | | | | | | | | | |
| --- | --- | --- | --- | --- | --- | --- | --- | --- | --- | --- | --- | --- |
| **Cases** | | **Homogeneity Correction** | | **Sum of Squares** | | **df** | | **Mean Square** | | **F** | | **p** |
| Genus |  | Welch |  | 0.701 |  | 9.000 |  | 0.078 |  | 133.420 |  | < .001 |
| Residual |  | Welch |  | 0.329 |  | 64.470 |  | 0.005 |  |  |  |  |
| *Note.*  Type III Sum of Squares | | | | | | | | | | | | |

| **Test for Equality of Variances (Levene's)** | | | | | | |
| --- | --- | --- | --- | --- | --- | --- |
| **F** | | **df1** | | **df2** | | **p** |
| 2.970 |  | 9.000 |  | 609.000 |  | 0.002 |

| **Games-Howell Post Hoc Comparisons - Genus** | | | | | | | | | | |
| --- | --- | --- | --- | --- | --- | --- | --- | --- | --- | --- |
|  | |  | | **Mean Difference** | | **SE** | | **t** | | **p** |
| Acanthochromis |  | Altrichthys |  | 0.046 |  | 0.005 |  | 9.289 |  | < .001 |
|  |  | Amblyglyphidodon |  | 0.054 |  | 0.003 |  | 16.758 |  | < .001 |
|  |  | Amphiprion |  | 0.087 |  | 0.003 |  | 33.127 |  | < .001 |
|  |  | Chromis |  | 0.019 |  | 0.005 |  | 4.140 |  | 0.009 |
|  |  | Dascyllus |  | 0.041 |  | 0.007 |  | 5.563 |  | < .001 |
|  |  | Hemiglyphidodon |  | 0.050 |  | 0.004 |  | 13.112 |  | < .001 |
|  |  | Lepidozygus |  | 0.044 |  | 0.004 |  | 10.489 |  | < .001 |
|  |  | Neoglyphidodon |  | 0.022 |  | 0.004 |  | 5.242 |  | < .001 |
|  |  | Pomacentrus |  | 0.017 |  | 0.007 |  | 2.424 |  | 0.452 |
| Altrichthys |  | Amblyglyphidodon |  | 0.008 |  | 0.005 |  | 1.696 |  | 0.789 |
|  |  | Amphiprion |  | 0.041 |  | 0.005 |  | 8.893 |  | < .001 |
|  |  | Chromis |  | -0.027 |  | 0.006 |  | -4.438 |  | 0.003 |
|  |  | Dascyllus |  | -0.005 |  | 0.008 |  | -0.591 |  | 1.000 |
|  |  | Hemiglyphidodon |  | 0.004 |  | 0.005 |  | 0.747 |  | 0.999 |
|  |  | Lepidozygus |  | -0.002 |  | 0.006 |  | -0.308 |  | 1.000 |
|  |  | Neoglyphidodon |  | -0.023 |  | 0.006 |  | -4.066 |  | 0.006 |
|  |  | Pomacentrus |  | -0.029 |  | 0.008 |  | -3.672 |  | 0.093 |
| Amblyglyphidodon |  | Amphiprion |  | 0.033 |  | 0.003 |  | 11.638 |  | < .001 |
|  |  | Chromis |  | -0.035 |  | 0.005 |  | -7.450 |  | < .001 |
|  |  | Dascyllus |  | -0.013 |  | 0.007 |  | -1.812 |  | 0.721 |
|  |  | Hemiglyphidodon |  | -0.004 |  | 0.004 |  | -1.141 |  | 0.979 |
|  |  | Lepidozygus |  | -0.010 |  | 0.004 |  | -2.382 |  | 0.370 |
|  |  | Neoglyphidodon |  | -0.032 |  | 0.004 |  | -7.265 |  | < .001 |
|  |  | Pomacentrus |  | -0.037 |  | 0.007 |  | -5.382 |  | 0.034 |
| Amphiprion |  | Chromis |  | -0.068 |  | 0.004 |  | -15.734 |  | < .001 |
|  |  | Dascyllus |  | -0.046 |  | 0.007 |  | -6.470 |  | < .001 |
|  |  | Hemiglyphidodon |  | -0.037 |  | 0.003 |  | -10.837 |  | < .001 |
|  |  | Lepidozygus |  | -0.043 |  | 0.004 |  | -11.126 |  | < .001 |
|  |  | Neoglyphidodon |  | -0.065 |  | 0.004 |  | -16.354 |  | < .001 |
|  |  | Pomacentrus |  | -0.070 |  | 0.007 |  | -10.497 |  | 0.003 |
| Chromis |  | Dascyllus |  | 0.022 |  | 0.008 |  | 2.684 |  | 0.231 |
|  |  | Hemiglyphidodon |  | 0.031 |  | 0.005 |  | 5.993 |  | < .001 |
|  |  | Lepidozygus |  | 0.025 |  | 0.005 |  | 4.591 |  | 0.002 |
|  |  | Neoglyphidodon |  | 0.003 |  | 0.005 |  | 0.600 |  | 1.000 |
|  |  | Pomacentrus |  | -0.002 |  | 0.008 |  | -0.305 |  | 1.000 |
| Dascyllus |  | Hemiglyphidodon |  | 0.009 |  | 0.008 |  | 1.165 |  | 0.971 |
|  |  | Lepidozygus |  | 0.003 |  | 0.008 |  | 0.399 |  | 1.000 |
|  |  | Neoglyphidodon |  | -0.018 |  | 0.008 |  | -2.333 |  | 0.403 |
|  |  | Pomacentrus |  | -0.024 |  | 0.010 |  | -2.513 |  | 0.338 |
| Hemiglyphidodon |  | Lepidozygus |  | -0.006 |  | 0.005 |  | -1.220 |  | 0.964 |
|  |  | Neoglyphidodon |  | -0.027 |  | 0.005 |  | -5.690 |  | < .001 |
|  |  | Pomacentrus |  | -0.033 |  | 0.007 |  | -4.558 |  | 0.052 |
| Lepidozygus |  | Neoglyphidodon |  | -0.022 |  | 0.005 |  | -4.208 |  | 0.004 |
|  |  | Pomacentrus |  | -0.027 |  | 0.007 |  | -3.650 |  | 0.116 |
| Neoglyphidodon |  | Pomacentrus |  | -0.006 |  | 0.007 |  | -0.751 |  | 0.997 |

Sum of Precaudal Vertebral Lengths / Sum of all Vertebral Lengths:

| **ANOVA - PCV Length / Vertebral Column Length** | | | | | | | | | | | | |
| --- | --- | --- | --- | --- | --- | --- | --- | --- | --- | --- | --- | --- |
| **Cases** | | **Homogeneity Correction** | | **Sum of Squares** | | **df** | | **Mean Square** | | **F** | | **p** |
| Genus |  | Welch |  | 0.215 |  | 9.000 |  | 0.024 |  | 113.049 |  | < .001 |
| Residual |  | Welch |  | 0.053 |  | 58.107 |  | 9.127e -4 |  |  |  |  |
| *Note.*  Type III Sum of Squares | | | | | | | | | | | | |

| **Test for Equality of Variances (Levene's)** | | | | | | |
| --- | --- | --- | --- | --- | --- | --- |
| **F** | | **df1** | | **df2** | | **p** |
| 1.477 |  | 9.000 |  | 260.000 |  | 0.156 |

| **Games-Howell Post Hoc Comparisons - Genus** | | | | | | | | | | |
| --- | --- | --- | --- | --- | --- | --- | --- | --- | --- | --- |
|  | |  | | **Mean Difference** | | **SE** | | **t** | | **p** |
| Acanthochromis |  | Altrichthys |  | 0.047 |  | 0.003 |  | 15.932 |  | < .001 |
|  |  | Amblyglyphidodon |  | 0.068 |  | 0.004 |  | 17.746 |  | < .001 |
|  |  | Amphiprion |  | 0.069 |  | 0.003 |  | 21.509 |  | < .001 |
|  |  | Chromis |  | 0.034 |  | 0.003 |  | 9.988 |  | < .001 |
|  |  | Dascyllus |  | 0.051 |  | 0.003 |  | 15.609 |  | < .001 |
|  |  | Hemiglyphidodon |  | 0.044 |  | 0.003 |  | 16.352 |  | < .001 |
|  |  | Lepidozygus |  | 0.083 |  | 0.003 |  | 28.780 |  | < .001 |
|  |  | Neoglyphidodon |  | 0.050 |  | 0.003 |  | 14.649 |  | < .001 |
|  |  | Pomacentrus |  | 0.057 |  | 0.003 |  | 17.563 |  | < .001 |
| Altrichthys |  | Amblyglyphidodon |  | 0.021 |  | 0.004 |  | 5.102 |  | < .001 |
|  |  | Amphiprion |  | 0.022 |  | 0.004 |  | 6.165 |  | < .001 |
|  |  | Chromis |  | -0.013 |  | 0.004 |  | -3.410 |  | 0.050 |
|  |  | Dascyllus |  | 0.004 |  | 0.004 |  | 1.197 |  | 0.967 |
|  |  | Hemiglyphidodon |  | -0.002 |  | 0.003 |  | -0.743 |  | 0.999 |
|  |  | Lepidozygus |  | 0.036 |  | 0.003 |  | 10.969 |  | < .001 |
|  |  | Neoglyphidodon |  | 0.003 |  | 0.004 |  | 0.752 |  | 0.999 |
|  |  | Pomacentrus |  | 0.010 |  | 0.004 |  | 2.770 |  | 0.254 |
| Amblyglyphidodon |  | Amphiprion |  | 9.028e -4 |  | 0.004 |  | 0.208 |  | 1.000 |
|  |  | Chromis |  | -0.034 |  | 0.004 |  | -7.571 |  | < .001 |
|  |  | Dascyllus |  | -0.017 |  | 0.004 |  | -3.823 |  | 0.014 |
|  |  | Hemiglyphidodon |  | -0.024 |  | 0.004 |  | -5.882 |  | < .001 |
|  |  | Lepidozygus |  | 0.015 |  | 0.004 |  | 3.654 |  | 0.023 |
|  |  | Neoglyphidodon |  | -0.018 |  | 0.004 |  | -4.100 |  | 0.005 |
|  |  | Pomacentrus |  | -0.011 |  | 0.004 |  | -2.562 |  | 0.298 |
| Amphiprion |  | Chromis |  | -0.035 |  | 0.004 |  | -8.792 |  | < .001 |
|  |  | Dascyllus |  | -0.018 |  | 0.004 |  | -4.583 |  | 0.002 |
|  |  | Hemiglyphidodon |  | -0.024 |  | 0.003 |  | -7.174 |  | < .001 |
|  |  | Lepidozygus |  | 0.014 |  | 0.004 |  | 3.993 |  | 0.011 |
|  |  | Neoglyphidodon |  | -0.019 |  | 0.004 |  | -4.868 |  | < .001 |
|  |  | Pomacentrus |  | -0.012 |  | 0.004 |  | -3.157 |  | 0.135 |
| Chromis |  | Dascyllus |  | 0.017 |  | 0.004 |  | 4.264 |  | 0.006 |
|  |  | Hemiglyphidodon |  | 0.010 |  | 0.004 |  | 2.916 |  | 0.144 |
|  |  | Lepidozygus |  | 0.049 |  | 0.004 |  | 13.187 |  | < .001 |
|  |  | Neoglyphidodon |  | 0.016 |  | 0.004 |  | 3.796 |  | 0.017 |
|  |  | Pomacentrus |  | 0.023 |  | 0.004 |  | 5.712 |  | 0.002 |
| Dascyllus |  | Hemiglyphidodon |  | -0.007 |  | 0.003 |  | -1.932 |  | 0.649 |
|  |  | Lepidozygus |  | 0.032 |  | 0.004 |  | 8.830 |  | < .001 |
|  |  | Neoglyphidodon |  | -0.002 |  | 0.004 |  | -0.386 |  | 1.000 |
|  |  | Pomacentrus |  | 0.006 |  | 0.004 |  | 1.447 |  | 0.891 |
| Hemiglyphidodon |  | Lepidozygus |  | 0.039 |  | 0.003 |  | 12.402 |  | < .001 |
|  |  | Neoglyphidodon |  | 0.005 |  | 0.004 |  | 1.443 |  | 0.907 |
|  |  | Pomacentrus |  | 0.012 |  | 0.003 |  | 3.595 |  | 0.087 |
| Lepidozygus |  | Neoglyphidodon |  | -0.033 |  | 0.004 |  | -9.013 |  | < .001 |
|  |  | Pomacentrus |  | -0.026 |  | 0.004 |  | -7.350 |  | < .001 |
| Neoglyphidodon |  | Pomacentrus |  | 0.007 |  | 0.004 |  | 1.803 |  | 0.725 |

Linear Regression of BCA/TBA ratio and Standard Length:

| **Model Summary** | | | | | | | | |
| --- | --- | --- | --- | --- | --- | --- | --- | --- |
| **Model** | | **R** | | **R²** | | **Adjusted R²** | | **RMSE** |
| 1 |  | 0.267 |  | 0.071 |  | 0.070 |  | 0.029 |

| **ANOVA** | | | | | | | | | | | | |
| --- | --- | --- | --- | --- | --- | --- | --- | --- | --- | --- | --- | --- |
| **Model** | |  | | **Sum of Squares** | | **df** | | **Mean Square** | | **F** | | **p** |
| 1 |  | Regression |  | 0.039 |  | 1 |  | 0.039 |  | 47.317 |  | < .001 |
|  |  | Residual |  | 0.507 |  | 617 |  | 8.224e -4 |  |  |  |  |
|  |  | Total |  | 0.546 |  | 618 |  |  |  |  |  |  |

| **Coefficients** | | | | | | | | | | | | |
| --- | --- | --- | --- | --- | --- | --- | --- | --- | --- | --- | --- | --- |
| **Model** | |  | | **Unstandardized** | | **Standard Error** | | **Standardized** | | **t** | | **p** |
| 1 |  | (Intercept) |  | 0.216 |  | 0.003 |  |  |  | 77.471 |  | < .001 |
|  |  | Standard Length (mm) |  | 2.891e -4 |  | 4.203e -5 |  | 0.267 |  | 6.879 |  | < .001 |

I’m including this in case anybody asks if the BCA/TBA ratio might be more reflective of different size classes in different genera vs actual differences between genera. This implies that standard length alone is a very poor predictor of BCA/TBA ratio. (Which is good!)

C. **Larval / Settler Swimming – U-crit**

Using the dataset presented in Fisher, R., Leis, J.M., Hogan, J.D. *et al.* Tropical larval and juvenile fish critical swimming speed (U-crit) and morphology data. *Sci Data* **9**, 45 (2022). <https://doi.org/10.1038/s41597-022-01146-3>)

| **ANOVA - ucrit** | | | | | | | | | | | | |
| --- | --- | --- | --- | --- | --- | --- | --- | --- | --- | --- | --- | --- |
| **Cases** | | **Homogeneity Correction** | | **Sum of Squares** | | **df** | | **Mean Square** | | **F** | | **p** |
| genus |  | Welch |  | 22535.453 |  | 10.000 |  | 2253.545 |  | 22.272 |  | < .001 |
| Residual |  | Welch |  | 64580.977 |  | 57.989 |  | 1113.675 |  |  |  |  |
| *Note.*  Type III Sum of Squares | | | | | | | | | | | | |

| **Test for Equality of Variances (Levene's)** | | | | | | |
| --- | --- | --- | --- | --- | --- | --- |
| **F** | | **df1** | | **df2** | | **p** |
| 7.759 |  | 10.000 |  | 377.000 |  | < .001 |

| **Games-Howell Post Hoc Comparisons - genus** | | | | | | | | | | |
| --- | --- | --- | --- | --- | --- | --- | --- | --- | --- | --- |
|  | |  | | **Mean Difference** | | **SE** | | **t** | | **p** |
| Abudefduf |  | Acanthochromis |  | 23.825 |  | 5.212 |  | 4.571 |  | 0.013 |
|  |  | Amblypomacentrus |  | 5.153 |  | 4.535 |  | 1.136 |  | 0.980 |
|  |  | Chromis |  | 2.606 |  | 4.314 |  | 0.604 |  | 1.000 |
|  |  | Chrysiptera |  | 12.829 |  | 3.217 |  | 3.988 |  | 0.009 |
|  |  | Dascyllus |  | 12.812 |  | 3.124 |  | 4.101 |  | 0.007 |
|  |  | Dischistodus |  | 8.959 |  | 3.659 |  | 2.449 |  | 0.356 |
|  |  | Neopomacentrus |  | -9.208 |  | 3.450 |  | -2.669 |  | 0.235 |
|  |  | Pomacentrus |  | -4.714 |  | 3.342 |  | -1.410 |  | 0.941 |
|  |  | Pristotis |  | -16.655 |  | 4.447 |  | -3.745 |  | 0.022 |
|  |  | Stegastes |  | 1.771 |  | 3.041 |  | 0.582 |  | 1.000 |
| Acanthochromis |  | Amblypomacentrus |  | -18.672 |  | 5.561 |  | -3.358 |  | 0.116 |
|  |  | Chromis |  | -21.219 |  | 5.382 |  | -3.943 |  | 0.038 |
|  |  | Chrysiptera |  | -10.996 |  | 4.550 |  | -2.417 |  | 0.447 |
|  |  | Dascyllus |  | -11.013 |  | 4.485 |  | -2.455 |  | 0.433 |
|  |  | Dischistodus |  | -14.866 |  | 4.872 |  | -3.051 |  | 0.192 |
|  |  | Neopomacentrus |  | -33.033 |  | 4.718 |  | -7.001 |  | 0.001 |
|  |  | Pomacentrus |  | -28.539 |  | 4.640 |  | -6.151 |  | 0.004 |
|  |  | Pristotis |  | -40.480 |  | 5.489 |  | -7.375 |  | < .001 |
|  |  | Stegastes |  | -22.054 |  | 4.428 |  | -4.981 |  | 0.024 |
| Amblypomacentrus |  | Chromis |  | -2.547 |  | 4.729 |  | -0.539 |  | 1.000 |
|  |  | Chrysiptera |  | 7.676 |  | 3.755 |  | 2.044 |  | 0.636 |
|  |  | Dascyllus |  | 7.659 |  | 3.677 |  | 2.083 |  | 0.619 |
|  |  | Dischistodus |  | 3.806 |  | 4.140 |  | 0.919 |  | 0.995 |
|  |  | Neopomacentrus |  | -14.361 |  | 3.957 |  | -3.629 |  | 0.108 |
|  |  | Pomacentrus |  | -9.866 |  | 3.864 |  | -2.554 |  | 0.396 |
|  |  | Pristotis |  | -21.807 |  | 4.851 |  | -4.496 |  | 0.014 |
|  |  | Stegastes |  | -3.382 |  | 3.606 |  | -0.938 |  | 0.991 |
| Chromis |  | Chrysiptera |  | 10.223 |  | 3.485 |  | 2.933 |  | 0.199 |
|  |  | Dascyllus |  | 10.206 |  | 3.400 |  | 3.002 |  | 0.185 |
|  |  | Dischistodus |  | 6.353 |  | 3.897 |  | 1.630 |  | 0.852 |
|  |  | Neopomacentrus |  | -11.814 |  | 3.702 |  | -3.191 |  | 0.115 |
|  |  | Pomacentrus |  | -7.320 |  | 3.601 |  | -2.032 |  | 0.632 |
|  |  | Pristotis |  | -19.261 |  | 4.645 |  | -4.147 |  | 0.012 |
|  |  | Stegastes |  | -0.835 |  | 3.324 |  | -0.251 |  | 1.000 |
| Chrysiptera |  | Dascyllus |  | -0.017 |  | 1.817 |  | -0.009 |  | 1.000 |
|  |  | Dischistodus |  | -3.870 |  | 2.631 |  | -1.471 |  | 0.918 |
|  |  | Neopomacentrus |  | -22.037 |  | 2.333 |  | -9.446 |  | < .001 |
|  |  | Pomacentrus |  | -17.543 |  | 2.170 |  | -8.084 |  | < .001 |
|  |  | Pristotis |  | -29.484 |  | 3.648 |  | -8.081 |  | < .001 |
|  |  | Stegastes |  | -11.058 |  | 1.670 |  | -6.623 |  | < .001 |
| Dascyllus |  | Dischistodus |  | -3.853 |  | 2.517 |  | -1.531 |  | 0.895 |
|  |  | Neopomacentrus |  | -22.020 |  | 2.204 |  | -9.992 |  | < .001 |
|  |  | Pomacentrus |  | -17.526 |  | 2.031 |  | -8.631 |  | < .001 |
|  |  | Pristotis |  | -29.467 |  | 3.567 |  | -8.260 |  | < .001 |
|  |  | Stegastes |  | -11.041 |  | 1.484 |  | -7.441 |  | < .001 |
| Dischistodus |  | Neopomacentrus |  | -18.167 |  | 2.912 |  | -6.239 |  | < .001 |
|  |  | Pomacentrus |  | -13.673 |  | 2.783 |  | -4.912 |  | < .001 |
|  |  | Pristotis |  | -25.614 |  | 4.043 |  | -6.335 |  | < .001 |
|  |  | Stegastes |  | -7.188 |  | 2.414 |  | -2.978 |  | 0.159 |
| Neopomacentrus |  | Pomacentrus |  | 4.494 |  | 2.503 |  | 1.795 |  | 0.781 |
|  |  | Pristotis |  | -7.447 |  | 3.856 |  | -1.931 |  | 0.693 |
|  |  | Stegastes |  | 10.979 |  | 2.084 |  | 5.267 |  | < .001 |
| Pomacentrus |  | Pristotis |  | -11.941 |  | 3.760 |  | -3.176 |  | 0.109 |
|  |  | Stegastes |  | 6.484 |  | 1.901 |  | 3.412 |  | 0.033 |
| Pristotis |  | Stegastes |  | 18.425 |  | 3.495 |  | 5.272 |  | 0.002 |
